# Whole-genome duplication increases genetic diversity and load in outcrossing *Arabidopsis*

**DOI:** 10.1101/2025.01.12.632621

**Authors:** Jakub Vlček, Tuomas Hämälä, Cristina Vives Cobo, Emma Curran, Gabriela Šrámková, Tanja Slotte, Roswitha Schmickl, Levi Yant, Filip Kolář

## Abstract

Genetic variation underpins evolutionary change, but accumulation of slightly deleterious mutations also increases mutation load. There are multiple factors affecting the extent of load such as population size and breeding system, yet other potential determinants remain unexplored. A common macromutation, whole-genome duplication (WGD) occurs broadly across Eukaryotes, yet we lack a clear understanding of how WGD impacts neutral and selective processes within a population. Using forward simulations and empirical analysis of 632 short- and 16 long-read sequenced individuals of *Arabidopsis arenosa* (23 diploid and 42 natural autotetraploid populations), we test for the effects of WGD on genome-wide diversity and mutation load. Our simulations show how genetic variation gradually rises in autotetraploids due to increase of mutational target size. Moreover, mutation load increases due to relaxed purifying selection when deleterious mutations are masked by additional chromosome copies. Empirical data confirm these patterns, showing significant increase in nucleotide diversity, ratios of non-synonymous to synonymous SNPs, and number of indels and large structural variants in *A. arenosa* autotetraploids. However, a rather modest increase in load proxies together with a broad distribution and niche of autotetraploids suggests load accumulation has not (yet) limited their successful expansion. Overall, we demonstrate a complex interplay between neutral processes and purifying selection in shaping genetic variation following WGD and highlight ploidy as an important determinant of genetic diversity and mutation load in natural populations.

## Introduction

Genetic diversity is both a result and a determinant of evolutionary change. While some variation is recruited for adaptation, most mutations are at least mildly deleterious and often disrupt molecular networks that have been fine-tuned over the course of evolution (1, 2). The reduction in fitness of a population, particularly due to recurrent deleterious mutations, is referred to as mutation load (3–6). While DNA mutagenesis increases mutation load, purifying selection reduces it by removing deleterious alleles. Understanding the factors that influence the efficiency of purifying selection and thus the level of mutation load is key to assessment of the adaptive potential of natural populations and has direct relevance to environmental challenges (7), agriculture (8), conservation (9–11), and human health (12, 13).

Both theoretical and empirical studies have revealed several key determinants of mutation load in natural populations. Population size and the degree of inbreeding are particularly strong factors (6, 14, 15), whose effect in empirical data, however, differs depending on what load component is measured. Realized and masked load reflect whether fitness effect of deleterious alleles is expressed in current generation or hidden in heterozygous state (recessive alleles), respectively (6). Assuming deleterious mutations being recessive or partially recessive, masked load increases linearly with population size and realized load drops as deleterious mutations accumulate in heterozygous state (6). Inbreeding on the other hand unmasks recessive deleterious mutations, increasing realized load. Multiple empirical studies support the notion that various types of demographic changes modulate selection efficiency and mutation load (15–18). Breeding system is yet another factor that shapes levels of homozygosity and therefore the amount of genetic load (19, 20). Recombination rate shapes mutation load as higher recombination allows more efficient selection (21, 22). And finally, mutation load is determined by mutation rate itself (23). In sum, recent studies demonstrated several population genetic processes significantly affecting mutation load, yet we still miss a comprehensive view on the spectrum of determinants of mutation load in natural populations.

Whole genome duplication (WGD) is a macro-mutation that substantially affects genetic transmission and effective population size which in turn may shape genetic variation and mutation load. WGD has an essential role in the evolution of eukaryotes (24) and it is widespread in plants, particularly in crops (25, 26), yet its effect on different predictors of mutation load has not been addressed. Here we aim to fill in this gap by studying the direct effects of WGD in intraspecific polyploids, autopolyploids, where natural WGD has not been confounded by hybridization. Theory shows that polysomic inheritance typical for autopolyploids, where all homologous chromosomes recombine without pairing partner preferences, modulates several aspects of genetic transmission (27). After one round of WGD, the mutation rate per gene is expected to double (25, 28), while the rate of genetic drift is halved (29). Accordingly, genetic variation in terms of nucleotide diversity (π) is expected to double (4Neμ vs. 8Neμ) (27) if we assume population census size is equal. Purifying selection is thought to be weaker in polyploids, predicting an increase in the number and frequency of deleterious alleles (30). This is due to masking of recessive or partly recessive alleles by additional sets of chromosomes. In other words, their exposure to selection is less frequent due to decreased homozygosity in autotetraploids (q^2^ in diploids vs. q^4^ in autotetraploids).

Despite these clear expectations, empirical data on genetic diversity in autopolyploids are not straightforward. First, genetic diversity has only occasionally been found to be higher (but never doubled) in autotetraploids compared to diploids (31–33). Despite this, increased variation of evolutionarily constrained (genic) regions has been found in autotetraploids for single nucleotide polymorphisms (SNPs) (34, 35), transposable elements (TE) polymorphisms (36) and structural variants (SVs) (37) providing first indices that mutation load may accumulate more in tetraploid populations compared to their diploid counterparts. Nonetheless, estimating load directly from genetic diversity is challenging (6, 38) and we lack a truly integrative analysis of mutation load and its components over multiple types of genetic markers and load indices in a natural polyploid system. Here, we aim to fill the knowledge gap by using several mutation load indices that account for differences in demography between ploidy cytotypes and complementing them by population genetic simulations to set our empirical results in a wider context of non-equilibrium evolutionary states.

In order to deconstruct the impact of WGD on global variation and mutation load, we combine forward-in-time simulations with empirical analysis of range-wide SV, insertion/deletion (indel), and SNP diversity in a diploid-autotetraploid species, *Arabidopsis arenosa*. First, we use forward-in-time simulations parameterized by values guided by our dataset to generate relevant predictions for the dynamics of genomic variation and mutation load as a function of WGD. Then, we analyse 65 short-read sequenced populations and 16 long-read sequenced individuals to empirically test these predictions. *A. arenosa* encompasses diploids and established natural autotetraploids that were formed by a single WGD event between 19,000 to 30,000 generations ago (34, 39). Its outcrossing breeding system, large and demographically stable populations, and excellent background knowledge of ploidy distribution, niche preferences and evolutionary history allow for testing general hypotheses on the drivers of genomic variation in natural conditions (34, 39–45). In particular, we test the hypothesis that tetraploid populations accumulate more genetic variation globally and genetic load in particular. By addressing this hypothesis across SNPs, indels and SVs we provide robust evidence for the genome-wide effect of WGD on accumulation of genetic variation reflecting both increase in mutational target size and decrease of selection efficiency.

## Results

### Forward-in-time simulations show pervasive effects of WGD on genetic diversity and mutation load

By simulating diploid and autotetraploid (hereafter simply, ‘tetraploid’) populations of the same maximum population census size (carrying capacity ∼180,000 individuals, parameterized with our empirical data, see methods), we observed that neutral diversity (π_Neutral_) slowly increases after a diploid population at equilibrium goes through a WGD. However, the increase is gradual (Fig. 1A) and π_Neutral_ in tetraploids does not reach the expected double values of diploids within even half a million generations, i.e. more than an order of magnitude higher than the estimated age of *A. arenosa* tetraploids (Fig. S1). Although π_Neutral_ estimates are consistently higher without background selection (reduction of neutral diversity due to purging of linked deleterious variants), the trajectory is very similar when background selection is included (Fig. S1), suggesting that the overall increase in nucleotide diversity primarily reflects an increase in mutational target size after WGD (doubled number of chromosomes), rather than weakened purifying selection due to polysomic masking. We did not assume an initial bottleneck following the formation of the tetraploid lineage, as there were no traces of past population size change in *A. arenosa*, likely due to gradual circa- and post-WGD influx of additional variation from the founding diploid population in the area where the tetraploid originated (34, 39). Yet, even when we assumed such a bottleneck, the overall patterns were similar, although π_Neutral_ of tetraploids drops immediately after the WGD and reaches the value of diploids only after 30,000 simulated generations (Fig. S2).

**Figure 1:**
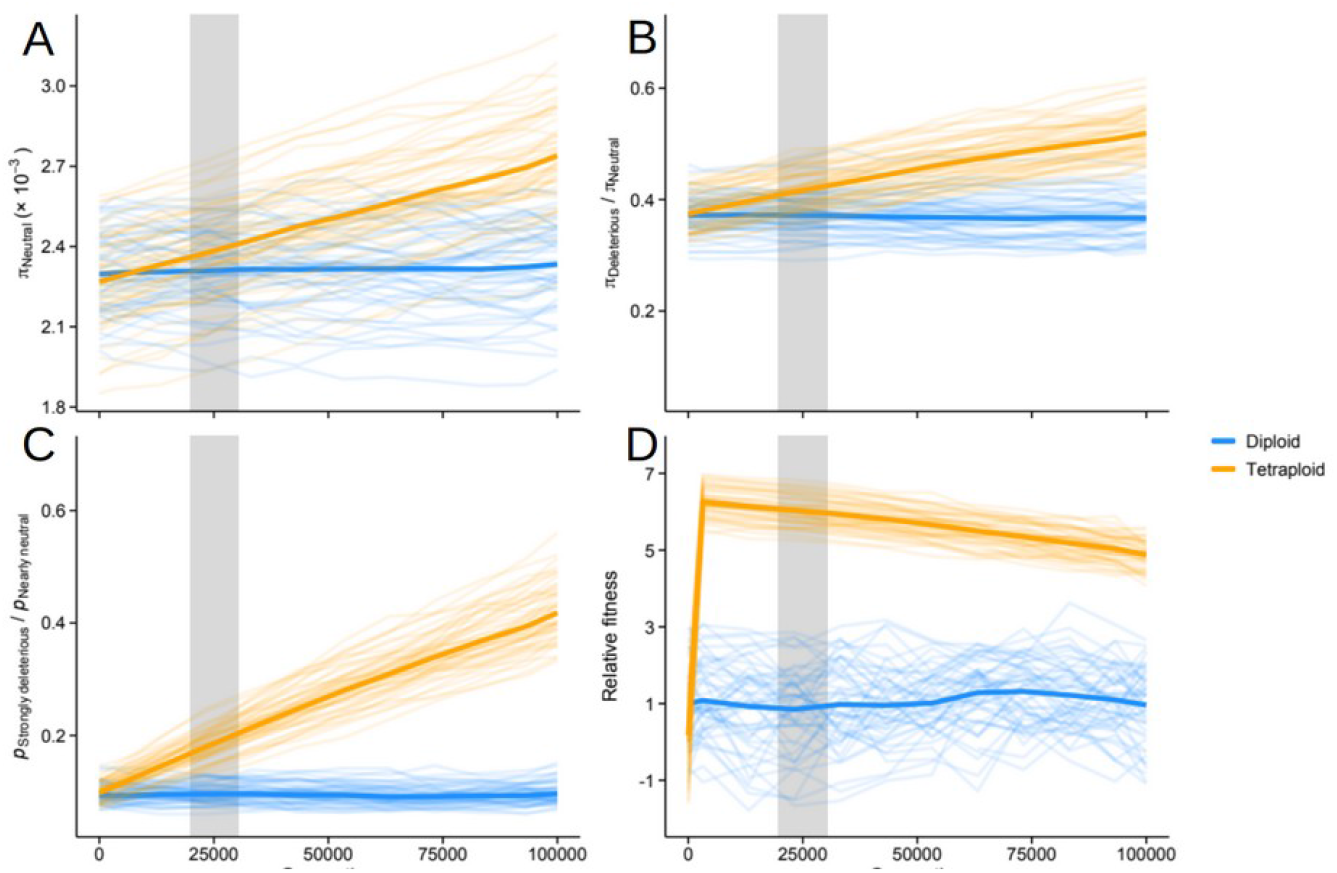
Forward simulation of genetic diversity and mutation load demonstrates pervasive effects of ploidy. Diploid (blue) and tetraploid (orange) populations during 100,000 generations following WGD. The carrying capacity of both cytotypes was the same and we assumed no founder bottleneck event in tetraploids (for bottleneck scenarios see supplementary figure S2). A) Neutral synonymous nucleotide diversity, B) ratio of non-synonymous to neutral nucleotide diversity (π_Deleterious_/π_Neutral_), C) ratio of strongly deleterious to nearly neutral polymorphisms D) average fitness of a population estimated as the product of fitness effects of all mutations. Fitness estimates were standardised based on values in diploids (mean = 1, SD = 1). In all panels, the solid lines show averages across 100 simulated repeats, while each of the simulation replicates are shown in transparent colours in the background. The grey area highlights the interval of estimated age of the extant *A. arenosa* tetraploids from Monnahan et al. (2019).

The accumulation of mutation load in tetraploids is also a gradual process. Two indices of mutation load–the ratio of deleterious to neutral nucleotide diversity and the ratio of strongly deleterious to nearly neutral polymorphisms–indicate that tetraploids accumulate a higher mutation load than diploids, with this difference becoming pronounced shortly after WGD (Fig. 1B & 1C). This result is robust to demographic effects as we see similar results if we assume a founding bottleneck at tetraploid formation (Fig. S2). Unsurprisingly, masking of deleterious mutations after WGD provides immediate fitness benefits for the nascent polyploids (Fig. 1D); interestingly however, these benefits are highly transient. As deleterious mutations accumulate, the fitness of the tetraploid population eventually falls below its diploid progenitor (Fig. S3).

In summary, our forward simulations show that WGD alters both neutral processes and the impact of purifying selection which should leave detectable footprints on genome-wide diversity in empirical datasets of diploid and autotetraploid populations.

### Genetic diversity and population structure in Arabidopsis arenosa

To decipher how WGD modulates genetic diversity and mutation load in natural populations, we assembled a range-wide sequencing dataset of 632 (222/410 diploid/tetraploid) *A. arenosa* individuals. We called genotypes in all accessions with available short-read sequencing data and included 126 newly sequenced individuals covering previously underrepresented regions/lineages resulting in 65 populations with at least 6 individuals sequenced (Fig. 2A, Tab. S1; average depth of coverage 24x). The structure of sampled populations corresponds with previous studies in the species (34, 39, 40, 42, 44): Neighbour-joining tree (Fig. 2D) and PCA (Fig. S4) show six distinct diploid lineages corresponding to those identified previously in a large range-wide sampling (40). Tetraploids form one dominant cluster, that is close to the ancestral (39) diploids from Western Carpathians, and three smaller clusters that show affinities to their sympatric diploid lineages, in line with the documented interploidy gene flow in these areas (34).

**Figure 2:**
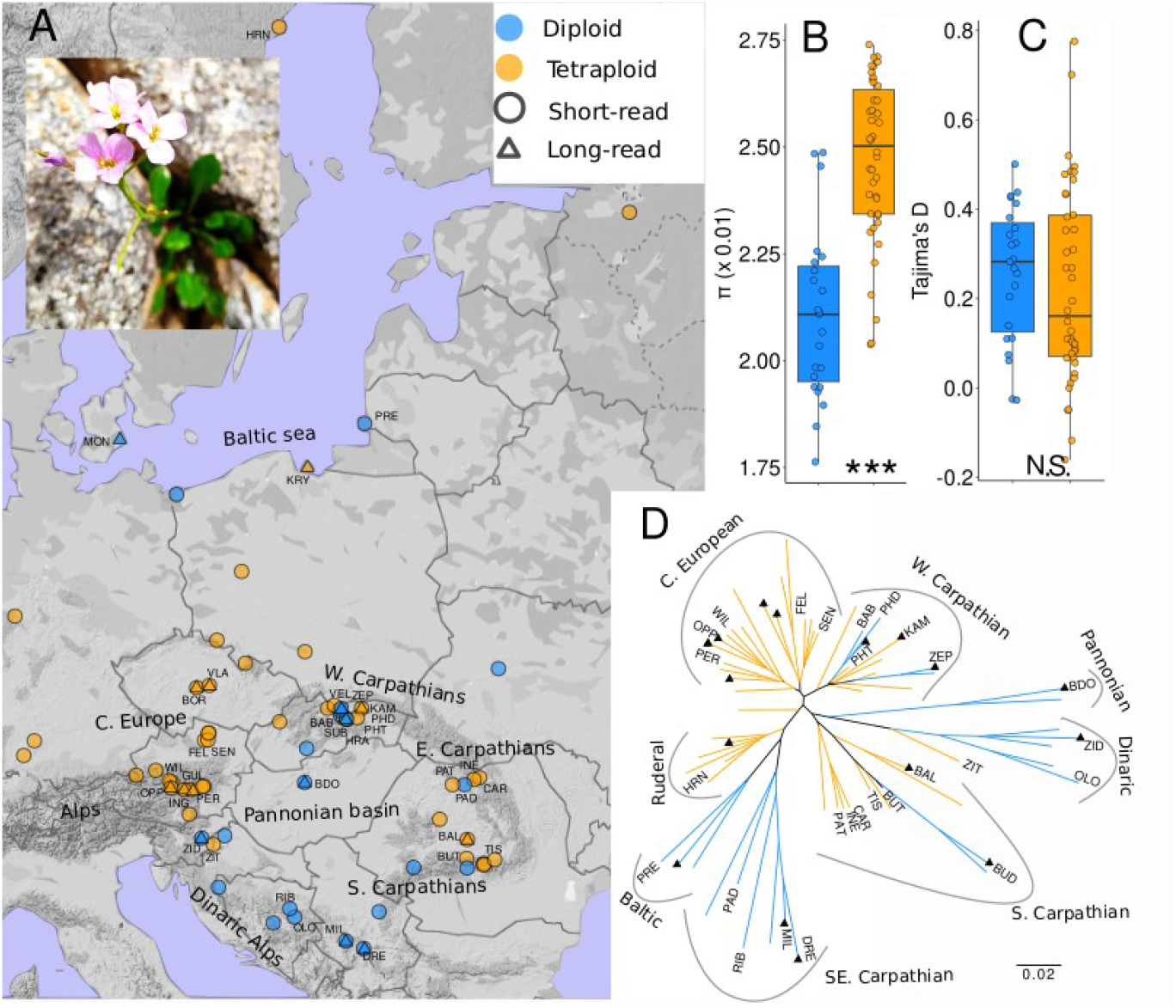
Range-wide sampling, genetic diversity, and structure of *Arabidopsis arenosa* populations. Diploid (blue) and tetraploid (orange) populations sampled throughout the species’ native range in Central and southeastern Europe: A) Locations of the full dataset of 65 populations sequenced for short reads (circles) and 16 individuals (triangles) sequenced for long reads. The photo shows flowering *A. arenosa* from W. Carpathians B,C) Within-population pairwise nucleotide diversity (π, B) and Tajima’s D (C) calculated based on four-fold degenerate SNPs in all populations (dots) downsampled to the same N of individuals (N=6). N.S. and asterisks indicate non-significant and significant difference between the ploidies as tested by Wilcoxon rank sum test, respectively. D) Genetic relationships of 65 populations depicted by a neighbor-joining tree based on Nei’s distances derived from 92,632 4-fold degenerate SNPs. Deeply-sequenced populations from the ‘core dataset’ are labelled with three letter code and populations with long read data are indicated by triangles.

Using populations as replicate observations, downsampled to the same number of individuals, we see that nucleotide diversity of putatively neutral 4-fold degenerate SNPs (π_s_) is 1.2 times higher in tetraploids (0.025) compared to diploids (0.021) on average (Fig. 2B, DF=62, β_ploidy_=3.9e^-3^,p_ploidy_<0.001, generalised linear model also accounting for an effect of sequencing depth, Table S2). The effect of ploidy on diversity is also robust to sampling, remaining statistically significant if we (i) sample the same number of chromosomes (not individuals) per population, (ii) exclude admixed tetraploid populations from interploidy contact zones (Table S2).

The observed difference in neutral diversity can be primarily attributed to the increase of population-scaled mutation rate in tetraploids (8Neμ) compared to diploids (4Neμ). Additional confounding influences of demographic processes within populations seem unlikely in *A. arenosa*, as we found no difference in Tajima’s D between ploidies (Fig. 2C; Wilcoxon rank-sum test of population means, W=544, p=0.409). Theoretically, increased mutation rate μ in tetraploids might also have contributed, but estimates of diploid and tetraploid germline mutation rate are not available for *A. arenosa*. Based on the insights from our simulations, we speculate that diversity in tetraploids is far less than double that in diploids (only 1.19x), because the relatively recent tetraploid cytotype has not yet reached mutation-drift equilibrium (Fig. 1A).

### Signatures of relaxed purifying selection in tetraploid A. arenosa in SNP data

We next examined whether relaxed purifying selection due to polysomic masking contributes to increased genetic variation upon WGD. To test for a difference in selection efficiency we compared selectively constrained 0-fold degenerate SNPs where any nucleotide change results in an amino acid replacement, and putatively neutral 4-fold degenerate SNPs where any nucleotide change is permitted without altering the encoded amino acid. To minimise the potential effects of sequencing errors, we focused on a subset of 27 deeply sequenced populations, hereafter called ‘core dataset’ (average sequencing depth of fourfold sites 35x for diploids and 38x for tetraploids (Fig. S5, S6), see Methods for details). Site frequency spectra (SFS) for both diploid and tetraploid populations showed an increased proportion of singleton 0-fold variants compared to 4-fold variants, which indicates that 0-fold sites are under purifying selection (Fig. 3 A & B). To test whether purifying selection is relaxed in tetraploids we used a ratio of overall 0-fold to 4-fold diversity (π_0_/π_4_) which serves as a proxy for the efficiency of selection relative to drift. This ratio is significantly higher in tetraploid populations compared to diploids (Fig. 3 C, Wilcox.: W=24, p=0.001), in line with the expectation of higher mutation load in tetraploid populations.

**Figure 3:**
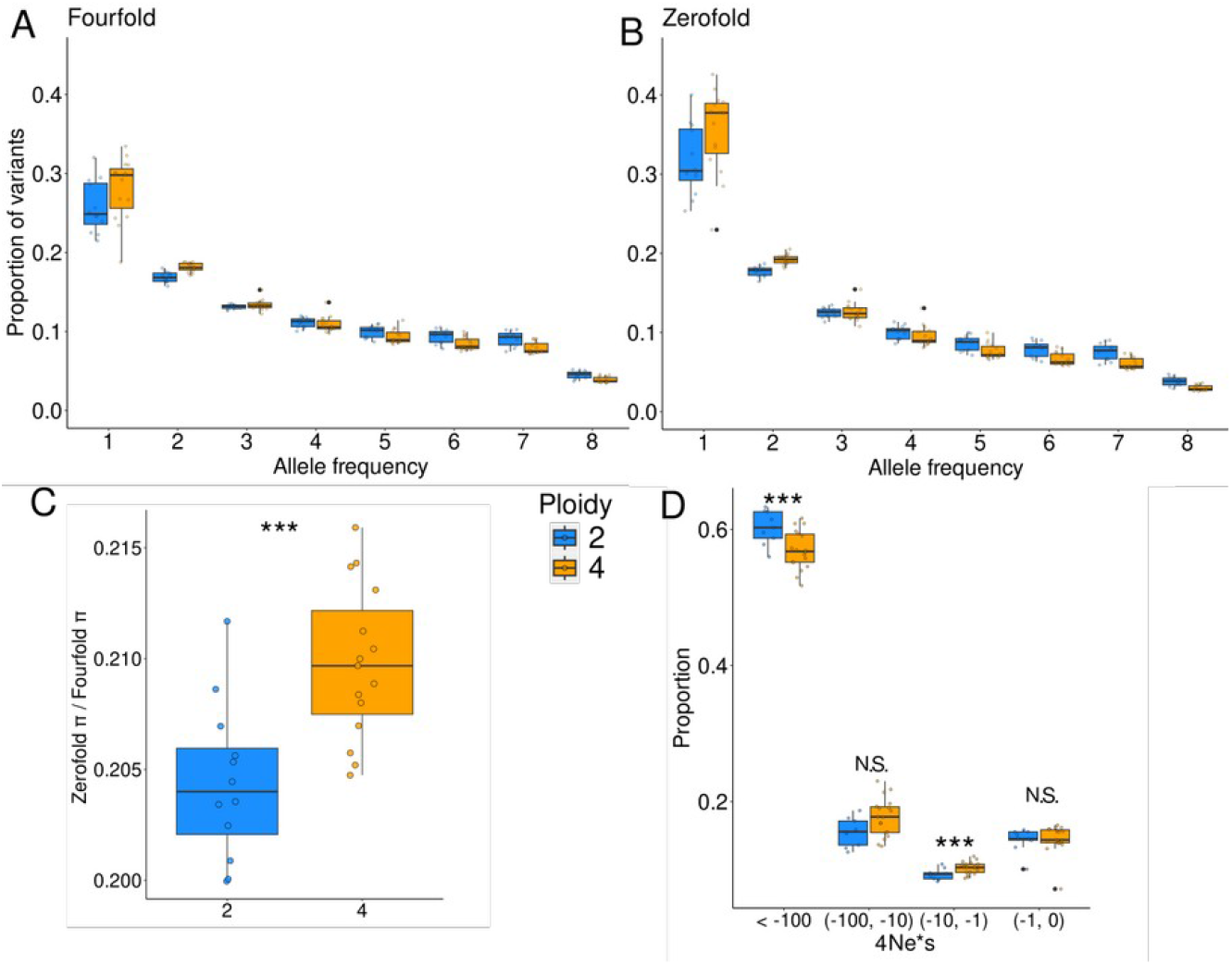
Signal of weaker purifying selection efficiency in tetraploid *A. arenosa* populations. Folded site frequency spectra (SFS) for A) putatively neutral 4-fold and B) putatively selected 0-fold SNP categories. The proportions are summarised over values inferred in our core dataset of 27 populations (dots), downsampled to the same N of allele copies. C) Ratio of 0-fold to 4-fold genetic diversity as a measure of selection efficiency in diploid and tetraploid populations in populations (dots) downsampled to the same N of individuals (N=6). D) Discretized distribution of deleterious fitness effects (DFE) estimated by polyDFE. A significant shift of DFE inferred in tetraploid populations towards moderately deleterious mutations likely reflects an effect of polysomic masking (see the main text). Boxplots indicate variation among the populations (dots), asterisks indicate statistically significant difference between cytotypes as tested by Wilcoxon rank-sum test.

**Figure 4:**
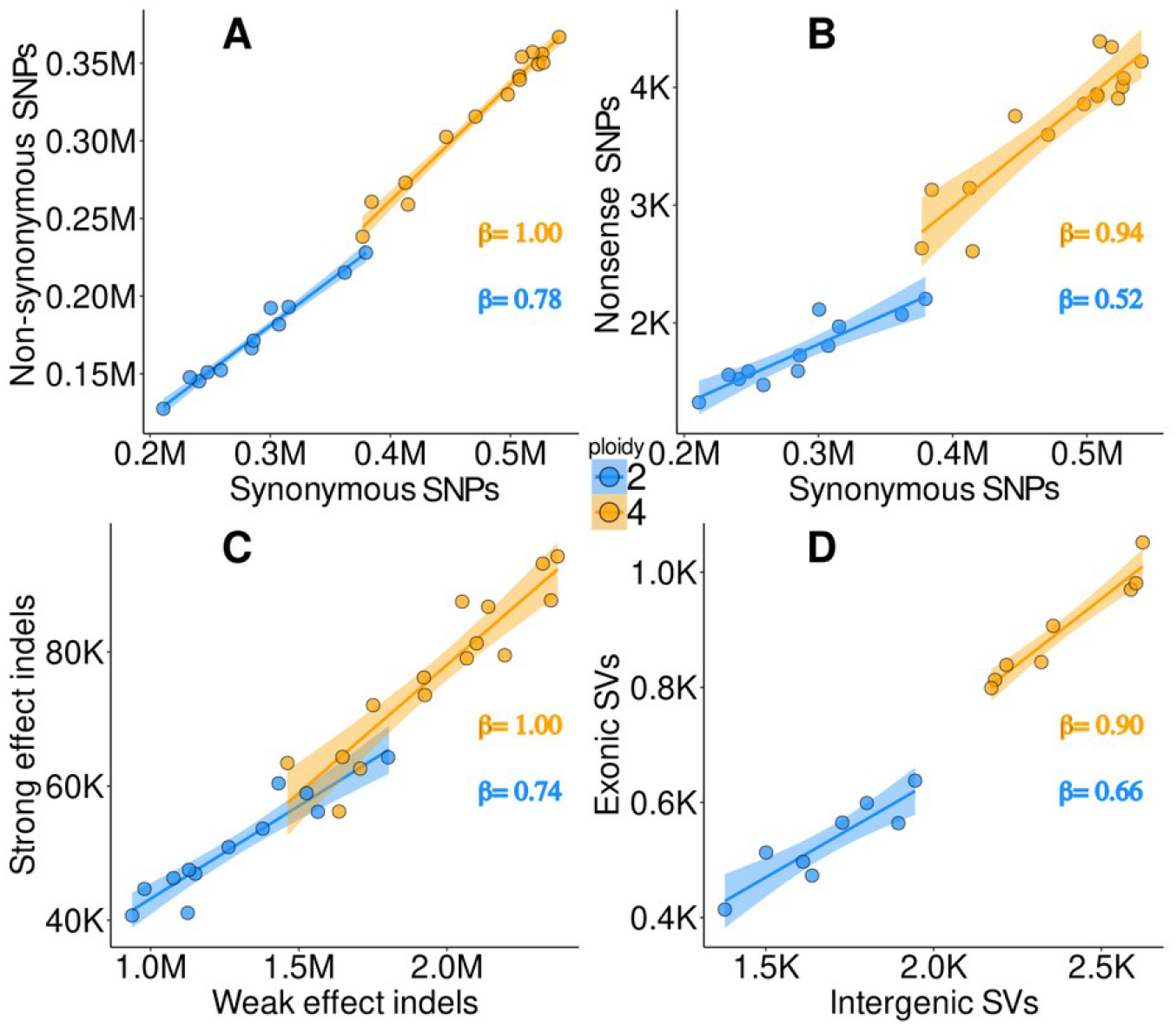
Accumulation of proportionally greater levels of deleterious variation in tetraploids across variant types. The relationship between the number of putatively neutral (horizontal axis) and deleterious (vertical axis) SNPs, insertion-deletion (indel) and structural variants (SVs) in diploid (blue) and tetraploid (orange) *A. arenosa* ‘core’ populations (A-C; downsampled to 6 individuals) or individuals (D). Tetraploids show not only consistently more variant sites of both categories, but also proportionally higher accumulation of deleterious variants (slope, β), an indicator of weakened efficiency of purifying selection (the interaction between N of putatively neutral variants and ploidy is significant for all variant types except for SVs, Table S4). Note that β was calculated for scaled and normalised data to allow direct comparison between site categories. A) Number of non-synonymous derived SNPs as a function of number of synonymous derived SNPs per population. B) Number of nonsense derived SNPs (inserting premature stop codons) as a function of number of synonymous derived SNPs per population. C) Number of strong effect derived indels as a function of number of weak effect derived indels per population. D) Total number of exonic SVs as a function of total number of intergenic SVs per individual. Fully annotated figures with population codes are available in supplementary data (Fig. S7).

We then asked whether the cytotypes differ in their distribution of fitness effects of new mutations (DFE). The DFE is shaped primarily by life history traits of a species (46), but as WGD leads to major changes in the dominance mode (28), it could affect DFE via modulation of selection efficiency. We inferred deleterious DFE from polymorphism in our core dataset by polyDFE (47). Surprisingly, we found that tetraploids showed a lower proportion of novel mutations with a likely highly deleterious effect. They also exhibited higher proportions of novel mutations with inferred moderately deleterious effects (Fig. 3D).

DFE inference is based on fitting a model that explains differences between 0-fold and putatively neutral 4-fold SFS within a population (48–50). Because the model assumes additivity, it does not account for the fact that, in tetraploids, putatively deleterious 0-fold variation may reach higher frequencies due to polysomic masking. This pattern in the SFS likely results in an apparent less deleterious DFE in tetraploids even if the real DFE is the same across ploidies. This interpretation is supported by the fact that DFE rarely changes within an outcrossing species (51, 52), however, at this stage we cannot conclusively differentiate between the effect of polysomic masking on DFE inference and ploidy-related change in actual DFE.

### Consistent signatures of relaxed purifying selection in tetraploids across all variant types

Finally, we explored whether the relaxed purifying selection suggested for SNPs is also reflected in short indels and larger structural variants (SVs) in the core short-read dataset (indels) and all 16 long-read sequenced individuals (SVs). We estimated mutation load based on variant counts, which can be applied consistently across all these data types. Consistent with neutral nucleotide polymorphism values, the total number of SNPs, indels and SVs was significantly higher in tetraploid compared to diploid populations. This held true both when accounted for the same N of individuals or chromosomes (Table S3). We thus asked whether it is only due to doubling of theta (4Neμ vs. 8Neμ) or also due to modulation of selection efficiency in coding regions in tetraploids. To empirically test this, we compared putatively neutral and deleterious categories of each variant type. In all cases, we see a positive linear correlation between the number of deleterious variants and putatively neutral variants across populations (Fig. 3, Fig S7). However, the slope of this relationship is always steeper in tetraploid populations (Fig. 3), and this is documented by the significant effect of interaction of neutral variation and ploidy level on deleterious variation for the population-level sampled variants (i.e., SNPs and indels, p<0.001, Table S4). In other words, tetraploids accumulate proportionally greater levels of deleterious variation than diploids do, even when taking neutral variation into account.

This observation is in line with the expectation of weaker purifying selection in tetraploids due to polysomic masking. In the case of SNPs, we used two different deleterious categories: all non-synonymous SNPs (Fig. 3A) and a subset of putatively large-effect nonsense SNPs (Fig. 3B). The difference between diploid and tetraploid slopes is bigger for nonsense SNPs compared to non-synonymous SNPs (Fig. 3A,B), indicating that the effect of polysomic masking is more pronounced for mutations with stronger fitness effects. This corresponds with the fact that more deleterious mutations tend to be also more recessive in *Arabidopsis* (53) and thus more effectively masked in tetraploids.

Running these analyses with the same number of chromosomes (as opposed to individuals) per population yielded qualitatively identical results (Fig. S8, Table S4), indicating robustness of the result to sampling strategy. Qualitatively identical results were also obtained when we focused on the total number of alleles per population (instead of sites) which quantifies total mutation load (38) (Fig. S9). The amount of realised load, quantified as counts of homozygous genotypes, is higher in diploids (Fig. S10) in line with expected higher homozygosity in diploid populations under equilibrium.

## Discussion

Here we demonstrated that whole-genome duplication (WGD) significantly increases genetic diversity and amount of deleterious variation in natural populations across a range of mutation types in an outcrossing plant species. A key novel insight is that we demonstrated the combined effect of WGD on both neutral and selective microevolutionary processes across a diverse array of genetic markers and natural populations varying in size and natural environments occupied.

Three salient points emerge from these analyses. First, our forward simulations and empirical data demonstrate that the increase of genetic diversity following WGD is primarily driven by increase in mutational target size. In the case of one round of WGD, mutational target size doubles and this results in an accumulation of a twice higher polymorphism if the number of individuals of pre-WGD and post-WGD population remains constant (28). Forward-in-time simulations support this theoretical expectation, however we show that the process of accumulation of polymorphism is gradual and doubled levels of polymorphism are not reached even after half a million simulated generations. In empirical data we see on average only 1.2-fold increase of neutral genetic polymorphism in tetraploid *A. arenosa* populations. This is in line with gradual accumulation of polymorphism following WGD, however, in empirical data we can not directly distinguish the effect of increase in mutational target size and the effect of demography as easily as in simulations where population size was kept constant. Overall, this suggests that differences in population history together with the gradual process of reaching mutation-drift equilibrium set a limit to the expected doubling of genetic diversity after one round of WGD. Consequently, this may explain why similar diversity has often been reported in empirical comparisons of diploid and autotetraploid natural populations, especially in studies focusing on recently diverging diploid-autopolyploid systems (e.g., 54–56, but see e.g. 57).

Second, our analyses also suggest that WGD has an effect on selective forces on top of the effect on neutral variation. Specifically, polysomic masking in autotetraploid populations is expected to relax purifying selection, leading to an accumulation of deleterious mutations in tetraploids. Indeed, the relative increase of genetic diversity of constrained sites is evident from the higher ratio of non-synonymous to synonymous nucleotide diversity and the altered distribution of fitness effects in *A. arenosa* tetraploids. Moreover, we show signatures of increased autotetraploid mutation load across different types of genetic variants that well-supplements similar effects in TE insertions, previously reported in a subset of *A. arenosa* populations (36). A consistent, yet nonsignificant trend in structural variants calls for further investigations of SV diversity at population level - an approach that is becoming feasible with the advances in long-read sequencing (e.g, 58). Altogether, our results suggest a pervasive impact of polysomic masking across the genome.

Third, in spite of clear signatures of accumulation of deleterious variation, our simulations and empirical data suggest that the recently formed tetraploid *A. arenosa* lineage may still benefit from lower realised load, i.e. the initial fitness effect of masking. This is consistent with overall successful niche expansion of the autotetraploid *A. arenosa* (44, 59) and empirical fitness estimates showing the autotetraploids perform as well as their diploid relatives across diverse environments (60). Additionally, the increased diversity may also serve as a pool of (potentially) adaptive variation especially in periods of environmental turmoil (61). That polyploid *A. arenosa* populations adapt from a large pool of standing variation, but also occasional novel mutations, has been documented over different extreme environments (62, 63). Nevertheless, our simulations suggest these advantages may be transient and the accumulating deleterious variation will likely result in negative fitness consequences and increased mutation load as the tetraploid ages, unless evolutionary “rescue” through rediploidisation and purging takes place (64).

It is important to note that our study focuses on empirical results of a single natural WGD event within a diverse and strictly outcrossing species with large and stable populations (34, 39 Fig 2). From studies in diploids, it is known that selfing and/or bottlenecks may affect load (17, 65, 66) but the interaction of these factors with ploidy remain unexplored. In other cases, nascent polyploid lineages may be affected by non-equilibrium demographic histories, as polyploid establishment may be accompanied by initial population bottleneck (67), range expansions (68), interploidy introgression (64, 69) and strong selection to adapt to the novel polyploid state (70–74). Thus, empirical investigations of diverse autopolyploid species spanning a range of breeding systems and demographic histories is a rich matter for further studies.

Crucially, our simulations suggested that the age of polyploid lineages determines the extent of variation that has newly accumulated in the tetraploid lineage. Recently formed polyploids may thus exhibit similar amount of deleterious variation as their diploid progenitors, yet higher fitness because of the masking of recessive deleterious in heterozygous state. In turn, we found surprisingly little effect of post-WGD interploidy gene flow on patterns of genetic diversity, despite rampant interploidy admixture (34). This might reflect a broadly shared gene pool across outcrossing *Arabidopsis* species and populations (73, 75) and the impact of introgression may be stronger in other mixed-ploidy groups where ancestral and admixing diploid lineages are more diverged (e.g. *Betula*, 76). Finally, the pattern of diversity and load may strongly differ in allopolyploids with disomic inheritance, where the inherited load from distinct diploid progenitors seem to play a crucial role (77–80).

In summary, our study highlights the dual role of WGD in increasing genetic diversity and mutation load, that is driven by both neutral processes and relaxed purifying selection. Polyploidy has occurred among a wide range of Eukaryotes, hence these findings situate polyploidy among important determinants of population genetic variation. Understanding how WGD shapes genetic diversity and mutation load in natural populations can inform strategies for managing genetic resources, improving crop resilience, and conserving biodiversity.

## Methods

### Simulation methods

To study factors influencing genetic diversity and mutation load in autotetraploids, we conducted individual-based, forward-in-time simulations using SLiM 3 (81). Polyploids are not directly supported by SLiM, and our approach for simulating autotetraploids is based on a script provided in the SLiM-Extras repository (https://github.com/MesserLab/SLiM-Extras). The simulations were conducted as non-Wright-Fisher (nonWF) models, which allow relaxing many assumptions of the standard Wright-Fisher (WF) models. In brief, generations in a nonWF model may overlap and individuals can reproduce multiple times which is a case of short-lived perennial *A. arenosa* (82). Fitness governs individuals’ survival to the next generation, rather than determining the mating success as in a WF model. Census population size (*N*) is a product of reproduction and survival, resulting in *N* variation across generations. This dynamic is controlled by the carrying capacity of the environment (*K*), which is enforced by scaling the fitness of the population based on the relationship of *K* and *N*, resulting in exponential growth until *K* is reached In SLiM 3, nonWF models also provide more flexibility in defining reproduction and recombination than WF models, which we leveraged in simulating tetraploids. This was achieved by simulating two diploid subpopulations and using the SLiM’s reproduction callback to allow crossovers to happen between chromosomes housed in the different subpopulations. Selection only acted on a single subpopulation, but dominance was a product of all four chromosomes. More details are provided in the annotated SLiM scripts, available at: https://github.com/thamala/polysim Genomic parameters were based on empirical studies conducted on *Arabidopsis* species; whenever possible, we used parameters inferred for our focal species *A. arenosa* as well as its close relatives from the genus *Arabidopsis*. However, the nonWF models employed here, especially the model implemented for tetraploids, are computationally more intensive than standard WF models, and therefore we used a rescaling approach to reduce computation times (83). Population size (*N*), mutation rate (*μ*), recombination rate (*r*), and selection coefficients (*s*) were rescaled by a factor of 20 while retaining the same product of *Nμ, Nr*, and *Ns* as the unscaled data (below we list the rescaled parameters). Note that we used the same scaling parameter for both diploids and tetraploids, resulting in the number of haploid genomes being double in tetraploids for a given *N*. Furthermore, rather than attempting to simulate whole genomes, we focused on a single chromosome of 100 kb and repeated the simulations 100 times.

We used an empirical mutation rate estimate (*μ* = 1.39*10-7) from *A. thaliana* (84), and assumed a ratio of deleterious to neutral *μ* of 2.76, as inferred for *A. lyrata* coding sequence (85). For crossover rate, we used a genome-wide average estimated for *A. arenosa* (*r* = 5.6*10-7) (86). Based on our estimates of the DFE, *s* for deleterious mutations were drawn from a gamma distribution with a mean of –0.048 and a shape parameter alpha of 0.228 (average across the diploid *A. arenosa* populations). Using data from *A. lyrata*, Huber et al. (87) co-estimated the distribution of selection and dominance coefficients (*h*) of new mutations. Following their results, we defined dominance coefficients based on the *s* of each mutation:

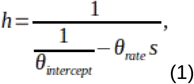

where *θ*_*intercept*_ = 0.978 and *θ*_*rute*_ = 50328, as inferred for *A. lyrata* (87). This continuous *h–s* model aims to capture the inverse relationship between selection and dominance often observed in empirical data (88, 89). Following Layman and Busch (90), we defined the fitness effect of each mutation as 1 + *h*_*x*_*S*, where *h*_*x*_ is a ploidy-independent dominance weight:

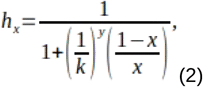

where *x* is the fraction of mutant copies (e.g., *x* = 0.5 for Aa and AAaa genotypes). Other components of the function (*k* and *y*) were solved as:

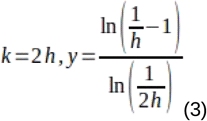

To define population sizes for the simulations, we transformed our empirical estimates of the population mutation rate (4*Neμ* in diploids and 8*Neμ* in tetraploids) to *Ne* by assuming *μ* = 6.95*10-9 (84), which gave an average *Ne* estimate of 183,680 for diploids and 104,680 for tetraploids. However, as we were primarily interested in assessing the effects of WGD (rather than *Ne*), we assumed the same *N* for both cytotypes. We started by establishing a diploid population with *K* = 9184 (corresponding to unscaled *N* of 183,680). After conducting a burn-in of 10*K* generations, an autotetraploid population was founded by duplicating the genomes of either 10 individuals (bottleneck scenario) or the whole population (non-bottleneck scenario presented in Fig. 1). We acknowledge that switching the whole population to polyploidy is likely unrealistic, but it allows us to distinguish the effects of WGD from the founding bottleneck and it also approaches a realistic situation for *A. arenosa* (34, 39) when the diversity of nascent autopolyploid population is enriched by post-WGD unidirectional gene flow from sympatric progenitor diploid population via unreduced gametes (69). To evaluate the mutation load of the newly founded autotetraploid population, we estimated nucleotide diversity for deleterious (π_Deleterious_) and neutral (π_Neutral_) variants, calculated the number of strongly deleterious (*s* < –0.01) and nearly neutral (*s* > –0.0001) mutations, and estimated fitness for each individual as a product of fitness effects of all mutations. These estimates were then compared to populations that remained diploid throughout the simulation.

### Empirical data sampling and sequencing

We gathered whole-genome short-read sequencing data of *A. arenosa* from all available published genome-sequencing datasets including populations with at least 6 and more well-covered individuals (34, 62, 63, 75). We complemented them with additional sequencing (126 individuals, 13 populations) to representatively cover all *A. arenosa* diploid and tetraploid lineages. In total, we gathered short read genome sequences for 634 accessions from 65 populations of *A. arenosa*. We removed reads of low quality and adaptor sequences with trimmomatic-0.36 (91) and we mapped refined reads to the new *A. arenosa* genome (92) by bwa-0.7.15 (93) with default setting. We used picard-2.8.1 to mark duplicate reads and called genotypes with GATK (v.3.7). We used HaplotypeCaller to infer genotypes per individual with respect to its ploidy. Information about ploidy of each individual was adopted from previous studies (published data) or it was detected by flow cytometry following our standard protocol (40) for the newly sequenced individuals.

To filter reliably called SNP genotypes we followed GATK best practices (94) and filtering strategies established in our previous studies involving autopolyploid populations (34, 63). For the SNP-based indices of mutation load and an analysis of population structure we used biallelic SNPs that passed these filtering parameters (FS>60.0 || SOR>3 || MQ<40 || MQRankSum<-12.5 || QD<2.0 || ReadPosRankSum<-8.0). Further, we masked genes with excessive heterozygosity with at least five fixed heterozygous SNPs in at least two diploid populations as being potentially paralogous genes following Monnahan et al. (34). Also we masked sites with excessive depth of coverage defined as sites with depth that exceeded twice standard deviation of depth of coverage in at least 20 individuals. Retaining only populations with at least 6 individuals left us with a ‘full’ dataset of 65 populations (632 individuals) with 54,179,280 SNPs of average sequencing depth of 24.2.

For methods that depend on precise inference of allele frequency spectra, we used only populations with mean SNP depth above 23. Applying such criteria left us with 27 sufficiently deeply sequenced ‘core dataset’ populations (12 diploid and 15 tetraploid) with mean depth of coverage of 36.1 (Figure S5).

### Indel variant calling and filtration

For the indel analysis we focused on the 27 deeply short-read sequenced core populations. We considered only sites that passed GATK best practice filters with the same parameters as for SNPs (FS>40.0 || SOR>3 || MQ<40 || MQRankSum<-12.5 || QD<2.0 || ReadPosRankSum<-4.0 || INFO/DP>2*mean coverage’). On top of that we removed sites with an excess depth coverage calculated as two times the mean coverage of the whole dataset to avoid misassembled paralogous sites. Only indels supported by at least 5 reads per individual and with absolute length up to 20 bp were considered. We did not remove multiallelic indels due to their ubiquity (48% of sites were multiallelic); we rather split the overlapping indels and counted them as distinct variants. We recovered 14.4M reliable indels while removing 59.4% of the raw number of indels (35.5M) that were called by GATK.

### Structural variant detection, calling and filtration

Structural variants were recovered using Oxford Nanopore long-read sequencing Technology (ONT). We extracted high molecular weight DNA from A. arenosa leaves as described in Russo et al. 2022. The DNA concentration was checked on a Qubit Fluorometer 2.0 (Invitrogen) using the Qubit dsDNA HS Assay kit. Fragment sizes were assessed using the Genomic DNA Tapestation assay (Agilent). Removal of short DNA fragments and final purification to high molecular weight DNA was performed with the Circulomics Short Read Eliminator XS kit. ONT libraries were prepared using the Genomic DNA Ligation kit SQK-LSK109 following the manufacturer’s procedure. Libraries were loaded onto R9.4.1 PromethION Flow Cells and run on a PromethION Beta sequencer. Due to the rapid accumulation of blocked flow cell pores or due to apparent read length anomalies on some runs, flow cells used in the runs were treated with a nuclease flush to digest blocking DNA fragments before loading with fresh libraries according to the ONT Nuclease Flush protocol (version NFL_9076_v109_revD_08Oct2018). FAST5 sequences produced by PromethION sequencer were basecalled using the Guppy6 (https://community.nanoporetech.com) high accuracy basecalling model (dna_r9.4.1_450bps_hac.cfg) and the resulting FASTQ files quality filtered by the basecaller. For read depth and quality of each sample see Table S5.

Minimap2 (v2.22) (95) was used to map the Nanopore reads against the reference genome of A. arenosa (92) with default parameters. We used the following SV calling pipeline developed for autotetraploid Cochlearia officinalis and validated using simulated autotetraploid data (37). Structural variants were identified using Sniffles2 (v2.0.6)(96) run in germline mode, limiting the minimum supporting reads depending on coverage (--minsupport auto). We only kept insertions and deletions between 50 bp and 100 kb in length, as read-alignment-based methods are less accurate at detecting other types of SVs as well as very large SVs. The R package updog based on a genotype-likelihood approach for allele frequency estimation in polyploids (97) was then used to estimate allelic dosage for each individual and SV. The depth of reference-supporting reads and variant-supporting reads for the SVs were retrieved from VCF files generated by Sniffles2. The multidog function was used for SVs with a minimum of 10 supporting reads with “norm model” and setting the ploidy level as 2 and 4 for diploids and tetraploids, respectively.

### Population structure inference

Relationships between all populations (full SNP dataset) have been visualised using PCA and Neighbor joining tree to confirm that our data match the structure observed in the previous studies which focused on the population structure in depth (39, 41, 34, 44). For the inference we used a frequency of 92632 4-fold degenerate biallelic SNPs with minimum genotype depth 8x and maximum 25% missing genotypes per site. This representation of unlinked sites was retrieved by pruning the 4-fold sites with maximum 0.25 linkage disequilibrium coefficient (r^2^) in 80 kb windows along the genome. We also removed sites where minor allele frequency was lower than 0.1. First we used principal component analysis (PCA) calculated using glPCA from the adegenet package (98) to visualise the population structure in multidimensional space. Then we calculated Nei’s distances (99) using StAMPP (100) and constructed a neighbour joining tree with the program SplitsTree (101).

### SNP-based analyses of population diversity

We calculated genome-wide nucleotide diversity of putatively neutral 4-fold, constrained 0-fold degenerate sites and Tajima’s D per each population by program Scantools (63, github.com/mbohutinská/ScanTools_ProtEvol) in the full SNP dataset. We used all biallelic SNPs with minimum genotype depth 8x and maximum 25% missing genotypes per site. We tested by Wilcoxon’s rank sum test whether diploid and tetraploid populations statistically differ in nucleotide diversity, ratio of 0-fold to 4-fold diversity and Tajima’s D. In order to account for possible effect of technical variation on our inference, we further tested for the effect of ploidy and sequencing depth, as an additional predictor, on the difference in nucleotide diversity using a general linear model from package stats in R version 4.1.2 (102). To equalise the sampling effort we (i) downsampled each population to 6 individuals and (ii) downsampled tetraploid populations to 4 individuals and diploid populations to 8 individuals (equal number of sampled allelic copies on 16 chromosomes).

### Distribution of fitness effects of SNP mutations

To test whether distribution of fitness effects (DFE, 2) differs between ploidy levels we first generated carefully polarised unfolded site frequency spectra (SFS) for 0-fold and 4-fold SNPs for each core population using the program est-sfs (103). As this method depends on correct characterization of allele frequency spectra, we focused on a subset of 27 sufficiently deeply sequenced populations (‘core dataset’). Est-sfs program input consists of allele counts of focal species and several outgroups. Allele counts per population of *A. arenosa* were parsed by an in-house python script sampling 16 chromosomes per population. For the outgroup counts we used *A. thaliana* and *Capsella rubella* and we merged the homologous sites of outgroups and our focal *A. arenosa* by genomic alignment of all species. Discrete DFE was then inferred using the re-polarised 0-fold and 4-fold SFS as inputs for each population by the program poly-DFE (47). This program allows inference of DFE with multiple parameters and different prior distributions of fitness effects. In order to explore the parameter space we ran models for all three basic modes of distributions (only deleterious, deleterious + beneficial displaced gamma distribution and deleterious + beneficial exponential distribution). Each distribution model was further refined by including or excluding a nuisance parameter for demography and a parameter for polarisation error. We then employed a series of likelihood ratio tests to determine the best-fitting model for each population. For comparison across populations, we finally used a model incorporating a deleterious + beneficial exponential distribution, a nuisance parameter, and a polarisation error parameter. This model showed the highest likelihood in model comparisons across the majority of populations. We tested by Wilcoxon’s rank sum test whether diploid and tetraploid populations statistically differ in the proportion of SNPs in each fitness bin.

### Correlational approach to determine mutation load across different variant types

We quantified mutation load over SNPs, short indels (both in stringently filtered ‘core’ short-read dataset of 27 populations) and SVs (all 16 long-read sequenced individuals, one per population) using a measure that allows consistent comparison among these different genetic markers. We calculated correlation between count of putatively neutral variants and selectively constrained variants per population and used these values as an index of mutation load, following studies of human populations (38).

First, we annotated the variants based on their putative phenotypic effect to get the two categories (neutral and constrained). For indels and SNPs we used SnpEff (5.1) (104) to annotate the putative phenotypic effect of each variant. For indels we used variants annotated as “HIGH” effect (typically frame-shift mutations) as constrained and “LOW” as neutral (intronic and intergenic variants). In SNPs we used two constrained categories a) non-synonymous SNPs (i.e. changing the resulting amino acid) b) nonsense SNPs (i.e. causing a premature stop codon) and one neutral category of synonymous SNPs. For SVs we used intergenic SVs (> 5 kb away from genes) as neutral category and exonic SVs (i.e. those at least partially covering an exon) as constrained. Then we counted all variable sites with allele frequency lower than 0.5 (to include only putatively derived variants) per each category and population. The populations were downsampled to 6 individuals for the SNP and indel dataset to achieve equal sampling effort. Alternatively we also downsampled tetraploid populations to 4 individuals and diploid populations to 8 individuals to explore whether sampling the same number of chromosomes per ploidy yields consistent results. Further we also counted alleles of a given category which represents an alternative summary of total mutation load (38). Finally, we also estimated realised mutation load as the number of homozygous genotypes. It shall be noted, however, that comparison of this index between cytotypes is directly affected by overall lower expected homozygosity in tetraploids (q^2^ in diploid while q^4^ in tetraploid populations).

Finally we tested the effect of ploidy on the number of constrained variants, while accounting for a ‘baseline’ neutral variation of each population, using generalised linear model from package stats in R version 4.1.2 (102). The model involved the number of constrained variants as response variable that was explained by ploidy and number of neutral variants, their interaction and the depth of coverage per population as predictors (constrainedN∼ploidy*neutralN+DP). To test for the effect of the interaction between ploidy and number of neutral variants, we compared two hierarchical models using likelihood ratio test: (i) with fixed effect of each predictor vs. (ii) with additional ploidy:neutralN interaction. Because the dependent variable is counts we used a Poisson distribution to model residual variance of the model.

## Data Availability

Sequence data that support the findings of this study are deposited in the NCBI (https://www.ncbi.nlm.nih.gov/bioproject/) under BioProjects PRJNA929698, PRJNA284572, PRJNA484107, PRJNA592307 and PRJNA667586 (short read data) and PRJEB83985 (long read data).

## Acknowledgements and funding sources

We are grateful to the members of the Plant ecological genomics team in Prague for support and constructive discussions. Further we thank Audry Le Veve, Magalena Bohutínská, Patrick Meirmans and Christian Parisod for valuable comments on earlier versions of this manuscript. This work has been supported by the Charles University Research Centre program (UNCE/24/SCI/006 to JV) and by the Czech Science Foundation (22-29078K to FK). Computational resources were provided by the e-INFRA CZ project (ID:90254), supported by the Ministry of Education, Youth and Sports of the Czech Republic. Sequencing was performed by the Norwegian Sequencing Centre, University of Oslo.

