## Supplementary data for "Whole-genome duplication increases genetic diversity and load in outcrossing *Arabidopsis*"

**Table S1:** Details on all populations of *A.arenosa* analyzed in this study.

| Pop | Ploidy | Lat. | Lon. | Lineage[1] | Admixed[2] | Core dataset[2,3] | Markers available | 4d depth[4] | 4d π | 4d Taj.D |
|---|---|---|---|---|---|---|---|---|---|---|
| BAB | 2 | 49.044 | 20.181 | West. Carp. | | YES | SNP, indel | 38.9 | 0.025 | -0.025 |
| BAL | 4 | 45.602 | 24.623 | South. Carp. | Y | YES | SNP, indel, SV | 44.0 | 0.024 | 0.388 |
| BDO | 2 | 47.458 | 18.925 | Pannonian | | YES | SNP, indel, SV | 27.8 | 0.022 | 0.288 |
| BUD | 2 | 45.467 | 25.228 | South. Carp. | | YES | SNP, indel, SV | 38.0 | 0.022 | 0.139 |
| BUT | 4 | 45.465 | 25.228 | South. Carp. | Y | YES | SNP, indel | 33.5 | 0.027 | 0.069 |
| CAR | 4 | 47.576 | 25.077 | South. Carp. | Y | YES | SNP, indel | 45.9 | 0.027 | 0.109 |
| DRE | 2 | 43.400 | 20.791 | South. Carp. | | YES | SNP, indel, SV | 40.7 | 0.019 | 0.427 |
| FEL | 4 | 48.457 | 15.402 | Central. Eu. | | YES | SNP, indel | 30.2 | 0.027 | 0.010 |
| HRA | 4 | 49.007 | 20.286 | West. Carp. | | YES | SNP, indel | 45.4 | 0.026 | 0.031 |
| HRN | 4 | 62.600 | 18.030 | Ruderal | Y | YES | SNP, indel | 34.4 | 0.027 | 0.776 |
| INE | 4 | 47.527 | 24.881 | South. Carp. | Y | YES | SNP, indel | 48.9 | 0.027 | 0.000 |
| KAM | 4 | 49.211 | 20.928 | West. Carp. | | YES | SNP, indel, SV | 39.5 | 0.023 | 0.496 |
| MIL | 2 | 43.511 | 20.442 | South. Carp. | | YES | SNP, indel, SV | 38.3 | 0.021 | 0.320 |
| OLO | 2 | 44.125 | 18.571 | Dinaric | | YES | SNP, indel | 31.3 | 0.018 | 0.430 |
| PAD | 2 | 47.402 | 24.546 | South. Carp. | | YES | SNP, indel | 32.4 | 0.023 | 0.268 |
| PAT | 4 | 47.402 | 24.546 | South. Carp. | Y | YES | SNP, indel | 27.1 | 0.027 | 0.023 |
| PER | 4 | 47.355 | 15.337 | Alpine | | YES | SNP, indel | 27.3 | 0.026 | 0.173 |
| PHD | 2 | 48.962 | 20.402 | West. Carp. | | YES | SNP, indel | 29.9 | 0.025 | 0.061 |
| PHT | 4 | 48.953 | 20.419 | West. Carp. | | YES | SNP, indel | 34.7 | 0.026 | -0.117 |
| PRE | 2 | 55.378 | 21.032 | Baltic | | YES | SNP, indel | 19.8 | 0.020 | 0.438 |
| RIB | 2 | 44.342 | 18.405 | South. Carp. | | YES | SNP, indel | 41.4 | 0.019 | 0.343 |
| SEN | 4 | 48.466 | 15.525 | Central. Eu. | | YES | SNP, indel | 41.6 | 0.027 | -0.050 |
| TIS | 4 | 45.570 | 25.608 | South. Carp. | | YES | SNP, indel | 44.1 | 0.027 | 0.083 |
| WIL | 4 | 47.326 | 14.230 | Alpine | | YES | SNP, indel | 46.0 | 0.025 | 0.353 |
| ZEP | 2 | 49.207 | 20.215 | West. Carp. | | YES | SNP, indel | 37.7 | 0.022 | 0.111 |
| ZID | 2 | 46.107 | 15.306 | Dinaric | | YES | SNP, indel,SV | 27.9 | 0.018 | 0.283 |
| ZIT | 4 | 46.094 | 15.344 | Dinaric | | YES | SNP, indel | 30.6 | 0.023 | 0.435 |
| MON | 2 | 54.963 | 12.551 | Baltic | | only SV | SV | NA | NA | NA |
| KRY | 4 | 54.380 | 19.423 | Ruderal | | only SV | SV | NA | NA | NA |
| BOR | 4 | 49.684 | 15.133 | Central. Eu. | | only SV | SNP, indel, SV | 14.9 | 0.025 | 0.246 |
| GUL | 4 | 47.282 | 14.928 | Central. Eu. | | only SV | SNP, indel, SV | 19.9 | 0.026 | 0.269 |
| ING | 4 | 47.284 | 14.682 | Alpine | | only SV | SNP, indel, SV | 19.4 | 0.024 | 0.467 |

| | | | | | | | | | | |
|---|---|---|---|---|---|---|---|---|---|---|
| OPP | 4 | 47.464 | 14.240 | Alpine | | only SV | SNP, indel, SV | 17.4 | 0.025 | 0.383 |
| SUB | 2 | 48.960 | 20.383 | West. Carp. | | only SV | SNP, indel | 80.5 | 0.025 | -0.027 |
| VEL | 2 | 49.162 | 20.154 | West. Carp. | | only SV | SNP, indel, SV | 15.1 | 0.022 | 0.075 |
| VLA | 4 | 49.735 | 15.175 | Central. Eu. | | only SV | SNP, indel, SV | 18.5 | 0.027 | 0.104 |
| BEL | 2 | 46.162 | 16.115 | Dinaric | | NO | SNP, indel | 13.0 | 0.019 | 0.324 |
| BGS | 4 | 47.628 | 13.002 | Ruderal | Y | NO | SNP, indel | 19.3 | 0.026 | 0.086 |
| BIH | 2 | 44.882 | 15.899 | Dinaric | | NO | SNP, indel | 11.8 | 0.019 | 0.229 |
| DRA | 4 | 45.442 | 25.224 | South. Carp. | Y | NO | SNP, indel | 11.6 | 0.022 | 0.067 |
| FOJ | 2 | 43.975 | 17.824 | Dinaric | | NO | SNP, indel | 15.0 | 0.020 | 0.501 |
| FUG | 4 | 48.631 | 15.557 | Central. Eu. | | NO | SNP, indel | 17.3 | 0.026 | 0.127 |
| GOR | 2 | 44.265 | 21.543 | South. Carp. | | NO | SNP, indel | 14.4 | 0.020 | 0.413 |
| HNE | 2 | 48.267 | 19.000 | Pannonian | | NO | SNP, indel | 12.5 | 0.022 | 0.257 |
| HOC | 4 | 47.370 | 15.387 | Alpine | | NO | SNP, indel | 17.7 | 0.026 | 0.100 |
| KAS | 4 | 46.688 | 14.872 | Alpine | | NO | SNP, indel | 20.0 | 0.023 | 0.519 |
| KLE | 4 | 50.244 | 16.848 | Ruderal | Y | NO | SNP, indel | 14.6 | 0.025 | 0.269 |
| KOS | 4 | 47.747 | 13.690 | Alpine | | NO | SNP, indel | 13.4 | 0.023 | 0.110 |
| KOW | 4 | 50.763 | 15.844 | Ruderal | Y | NO | SNP, indel | 13.4 | 0.024 | 0.484 |
| KRM | 2 | 50.119 | 25.740 | Baltic | | NO | SNP, indel | 18.9 | 0.021 | 0.381 |
| LAC | 4 | 45.595 | 24.635 | South. Carp. | Y | NO | SNP, indel | 15.3 | 0.020 | 0.701 |
| MIA | 4 | 50.503 | 18.938 | Ruderal | Y | NO | SNP, indel | 13.0 | 0.023 | -0.161 |
| MIE | 2 | 53.921 | 53.921 | Baltic | | NO | SNP, indel | 14.1 | 0.020 | 0.358 |
| RFT | 4 | 48.101 | 9.050 | Central. Eu. | | NO | SNP, indel | 12.4 | 0.023 | 0.310 |
| RZA | 2 | 45.378 | 22.758 | South. Carp. | | NO | SNP, indel | 12.6 | 0.021 | 0.204 |
| SPI | 4 | 48.989 | 20.775 | West. Carp. | | NO | SNP, indel | 12.9 | 0.023 | 0.304 |
| STE | 4 | 52.280 | 16.709 | Ruderal | Y | NO | SNP, indel | 19.4 | 0.026 | 0.149 |
| STG | 4 | 48.630 | 15.543 | Central. Eu. | | NO | SNP, indel | 16.4 | 0.027 | -0.048 |
| SWA | 4 | 48.448 | 9.422 | Central. Eu. | | NO | SNP, indel | 11.9 | 0.020 | 0.483 |
| TKO | 4 | 49.205 | 19.735 | West. Carp. | | NO | SNP, indel | 18.1 | 0.024 | 0.056 |
| TRE | 4 | 48.894 | 18.045 | West. Carp. | | NO | SNP, indel | 21.8 | 0.024 | 0.433 |
| TZI | 4 | 46.567 | 23.674 | South. Carp. | Y | NO | SNP, indel | 11.7 | 0.024 | 0.077 |
| VID | 2 | 45.364 | 24.638 | South. Carp. | | NO | SNP, indel | 13.1 | 0.021 | 0.110 |
| VOR | 4 | 47.499 | 14.170 | Alpine | | NO | SNP, indel | 17.9 | 0.025 | 0.355 |
| VYR | 4 | 59.414 | 30.346 | Ruderal | Y | NO | SNP, indel | 21.1 | 0.024 | 0.478 |
| WUL | 4 | 51.308 | 8.487 | Ruderal | Y | NO | SNP, indel | 11.0 | 0.022 | 0.099 |
| ZAP | 4 | 49.278 | 19.967 | West. Carp. | | NO | SNP, indel | 11.0 | 0.021 | 0.195 |

[1]'Lineage' corresponds to major genetic groups inferred by previous studies (Kolar et al. 2016, Monnahan et al 2019) [2]'Admixed' stands for population from interploidy contact zones with previously identified high levels of interploidy gene flow from sympatric but genetically divergent diploid lineages. [3]'Core dataset' indicates whether the population was used in the detailed analysis of mutation load across marker types. [4] 4d stands for four-fold degenerate sites. Nucleotide diversity (π) and Tajima's D (Taj.D) were calculated based on downsampling each population to 6 individuals.

**Table S2**: Results of general linear models testing the effect of ploidy and depth of coverage on nucleotide diversity of four fold degenerate sites (π~ploidy+depth). We ran the same model for different subsets: a) populations from the core deeply sequenced dataset, subsampled to 6 individuals, b) populations from the full dataset, subsampled to 16 chromosomes, i.e. the same number of alleles sampled in each ploidy c) populations from the full dataset subsampled to 6 individuals excluding tetraploid populations from known mixed-ploidy contact zones (Monnahan et al. 2019, see Table S1 for their list) where gene flow from divergent diploids could increase nucleotide diversity d) populations from the full dataset subsampled to 6 individuals. In the table we further report residual degrees of freedom, slope estimate (β), its standard error (SE) and significance value (Pr).

| Dataset | Residual DF | β ploidy | SE ploidy | Pr ploidy | β depth | SE depth | Pr depth |
|---|---|---|---|---|---|---|---|
| Core dataset 6 individuals | 24 | 4.72E-03 | 7.55E-04 | 1.83E-06 | 1.82E-05 | 5.23E-05 | 0.731 |
| Full dataset 16 chromosomes | 59 | 3.92E-03 | 5.08E-03 | 3.73E-03 | -1.74E-04 | 6.63E-05 | 0.011 |
| Full dataset 6 individuals without admixed tetraploids | 44 | 3.74E-03 | 4.87E-04 | 1.09E-09 | 6.04E-05 | 1.88E-05 | 0.0025 |
| Full dataset 6 individuals | 62 | 3.91E-03 | 4.38E-04 | 9.98E-13 | 7.20E-05 | 1.57E-05 | < 0.0001 |

**Table S3:** Results of Wilcoxon rank sum test, assessing the difference in total number of variable sites of each marker type between diploid and tetraploid populations in the core dataset of deeply sequenced populations. For SNPs and indels the tests were done for two different sampling strategies. First strategy was to take 6 individuals per each population corresponding to 12 and 24 chromosomes for diploids and tetraploids respectively. Second strategy was to take 4 tetraploid individuals and 8 diploids which corresponds to 16 chromosomes per each population for both cytotypes.

| Marker | Sampling | W | p | Diploid avg. | Tetraploid avg. |
|---|---|---|---|---|---|
| SNP | 6 individuals | 0 | < 0.0001 | 1,307,548 | 1,968,926 |
| SNP | 16 chromosomes | 11 | < 0.0001 | 1,399,362 | 1,762,645 |
| Indel | 6 individuals | 8 | < 0.0001 | 1,729,655 | 2,420,991 |
| Indel | 16 chromosomes | 41 | 0.016 | 1,865,910 | 2,147,939 |
| SV | per individual | 0 | < 0.0001 | 27,395 | 42,008 |

**Table S4:** Results of generalised linear models with Poisson distribution of errors testing the effect of ploidy level on the number of deleterious categories of SNPs, indels and structural variants while taking into account the amount of putatively neutral variants and sequencing depth (dp). In particular, we tested significance of the interaction of the number of neutral variants and ploidy, i.e. whether there is a differential proportional accumulation of deleterious categories between cytotypes. The statistical significance of the interaction was tested by likelihood ratio test comparing (i) a null model with only a direct effect of each factor (reference 'neutral' variant counts and ploidy) with (ii) a model also including their interaction. Two sampling strategies were used for SNPs and indels: 6 individuals and 16 chromosomes per population to normalise the population counts. Estimates of slope (β) and standard error (SE) are shown.

| Model equation | Dataset | Residual DF | Ploidy (β\| SE\| Pr) | Neutral variant (β\| SE\| Pr) | Depth (β\| SE\| Pr) | neutral:ploidy interaction (β\| SE\| Pr) | Deviance | Pr(>Chi) |
|---|---|---|---|---|---|---|---|---|
| Strong effect indels ~ (weak effect indels)*ploidy+dp | 6 ind. | 22 | 0.124\| 9.84E-03\| <0 | 4.94E-07\| 5.32E-09\| <0 | 1.89E-03\| 1.21E-04\| <0 | -2.89E-08\| 6.02E-09\| 1.59E-06 | 23 | < 0.0001 |
| Strong effect indels ~ (weak effect indels)*ploidy+dp | 16 chrom. | 22 | -0.121\| 0.011\| <0 | 4.69E-07\| 4.27E-09\| <0 | 1.73E-03\|1.30E-04\| <0 | 6.57E-08\| 6.42E-09\| <0 | 104.8 | < 0.0001 |
| Non-synonymous SNPs ~ (synonymous SNPs)*ploidy+dp | 6 ind. | 22 | 3.72E-01\| 6.01E-03\| <0 | 3.25E-06\| 1.39E-08\| <0 | 1.13E-03\| 4.92E-05\| <0 | 7.89E-07\| 1.65E-08\| <0 | 2274.5 | < 0.0001 |
| Non-synonymous SNPs ~ (synonymous SNPs)*ploidy+dp | 16 chrom. | 22 | 0.156\| 6.35E-3\| <0 | 3.01E-06\| 1.12E-08\| <0 | 1.09E-03\| 5.10E-05\| <0 | -3.37E-07\|1.64E-08\| <0 | 419.6 | < 0.0001 |
| Nonsense SNPs ~ (synonymous SNPs)*ploidy+dp | 6 ind. | 22 | 4.54E-03\| 4.44E-04\| 0.389 | 1.81E-06\| 1.29E-07\| <0 | 4.54E-03\| 4.44E-04\| <0 | 5.24E-07\| 1.51E-07\| 5.35E-04 | 12.01 | 0.0005 |
| Nonsense SNPs ~ (synonymous SNPs)*ploidy+dp | 16 chrom | 22 | -7.46E-02\| 5.77E-02\| 0.196 | 1.88E-06\| 1.04e-07\| <0 | 4.16E-07\| 4.66E-04\| <0 | 4.47E-07\| 1.50E-07\| 2.92E-3 | 8.88 | 0.0029 |
| Exonic SVs ~ (intron SVs)*ploidy | per ind. | 12 | 0.411\| 0.214\| 0.055 | 6.41E-04\| 8.57E-05\| <0 | NA | -1.38E-04\| 1.07E-04\| 0.197 | 1.66 | 0.197 |

**Table S5**: Read depth and quality of long read sequencing of each accession.

| Source population | Ploidy | Lineage | Depth | Read quality |
|---|---|---|---|---|
| BDO | 2x | Pannonian | 74 | 13.7 |
| ZID | 2x | Dinaric | 63 | 13.8 |
| BUD | 2x | South. Carp. | 63 | 14.1 |
| VEL | 2x | West. Carp. | 58 | 13.8 |
| SUB | 2x | West. Carp. | 61 | 13.8 |
| KAM | 4x | West. Carp. | 61 | 13.4 |
| GUL | 4x | Alpine | 267 | 14.3 |
| ING | 4x | Alpine | 41 | 13.5 |
| VLA | 4x | Central. Eu. | 74 | 13.3 |
| BAL | 4x | South. Carp. | 89 | 13.5 |
| BOR | 4x | Central. Eu. | 58 | 14.2 |
| OPP | 4x | Alpine | 75 | 14 |
| MON | 2x | Baltic | 33 | 14.3 |
| KRY | 4x | Ruderal | 64 | 13.8 |
| MIL | 2x | South. Carp. | 32 | 14.3 |
| DRE | 2x | South. Carp. | 29 | 14.2 |

### Supplementary figures

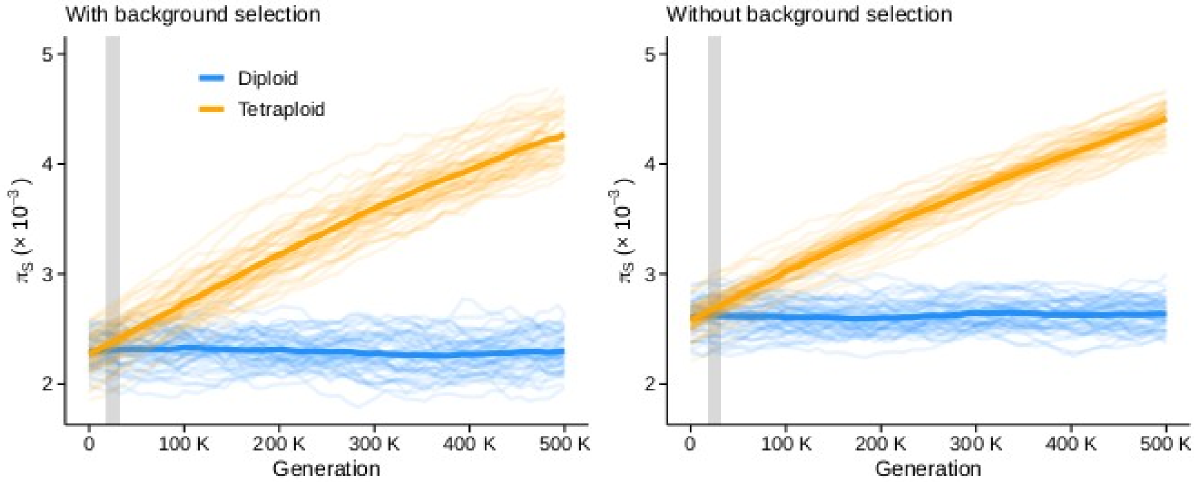


**Figure S1:** Simulated synonymous (neutral) nucleotide diversity over 500 000 generations after WGD in models assuming background selection (left) and under a simpler model without the effect of background selection (right). The solid lines show averages across 100 simulated repeats, while each of the simulation replicates are shown in semi-transparent colours in the background. Grey area highlights the interval of estimated age of the extant *A. arenosa* tetraploids following Monnahan et al. (2019).

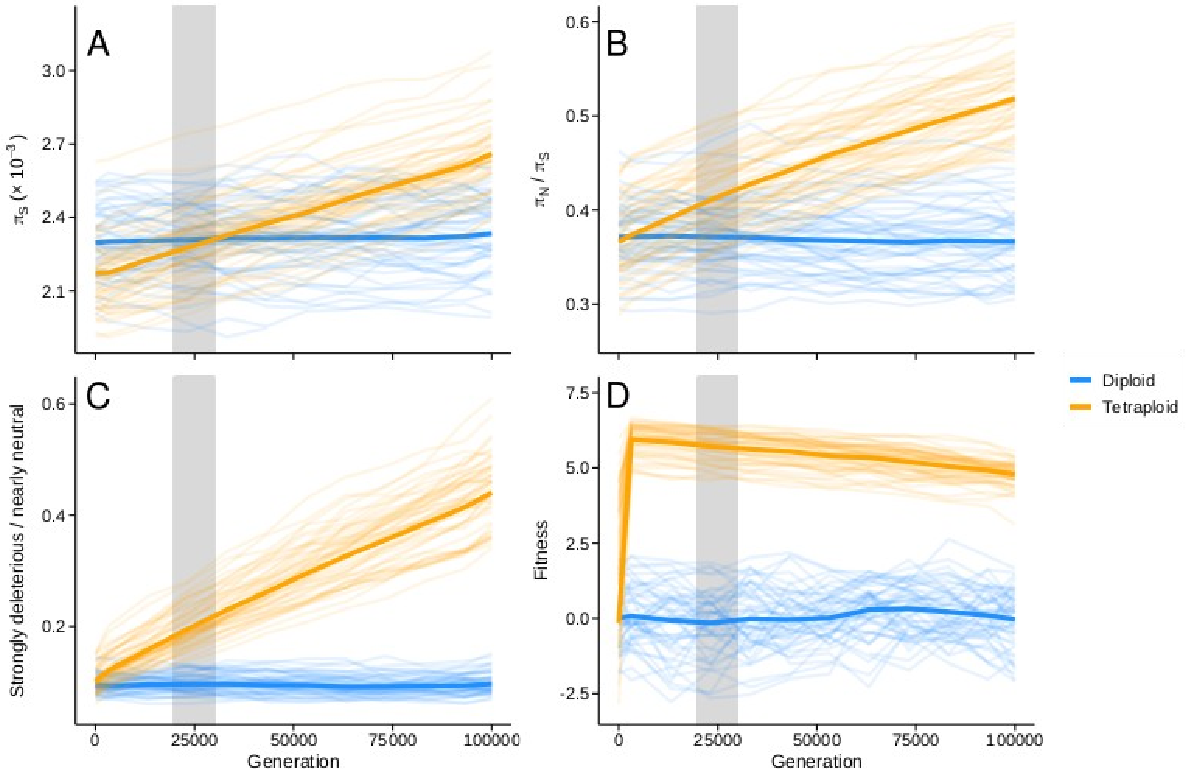


**Figure S2:** Forward simulation of genetic diversity and genetic load for diploid (blue) and tetraploid (orange) populations during 100 000 generations since WGD. At time zero, autotetraploid population is formed from a diploid population and experiences a bottleneck of 10 individuals. A) Neutral synonymous nucleotide diversity B) Ratio of non-synonymous to synonymous nucleotide diversity, C) Ratio of strongly deleterious to nearly neutral variants D) Average fitness of a population estimated as the product of fitness effects of all mutations. The fitness estimates were standardised based on values in diploids (mean = 0, SD = 1). In all panels, the solid lines show averages across 100 simulated repeats, while each of the simulation replicates are shown in transparent colours in the background. Grey area highlights the interval of estimated age of the extant *A. arenosa* tetraploids following Monnahan et al. (2019).

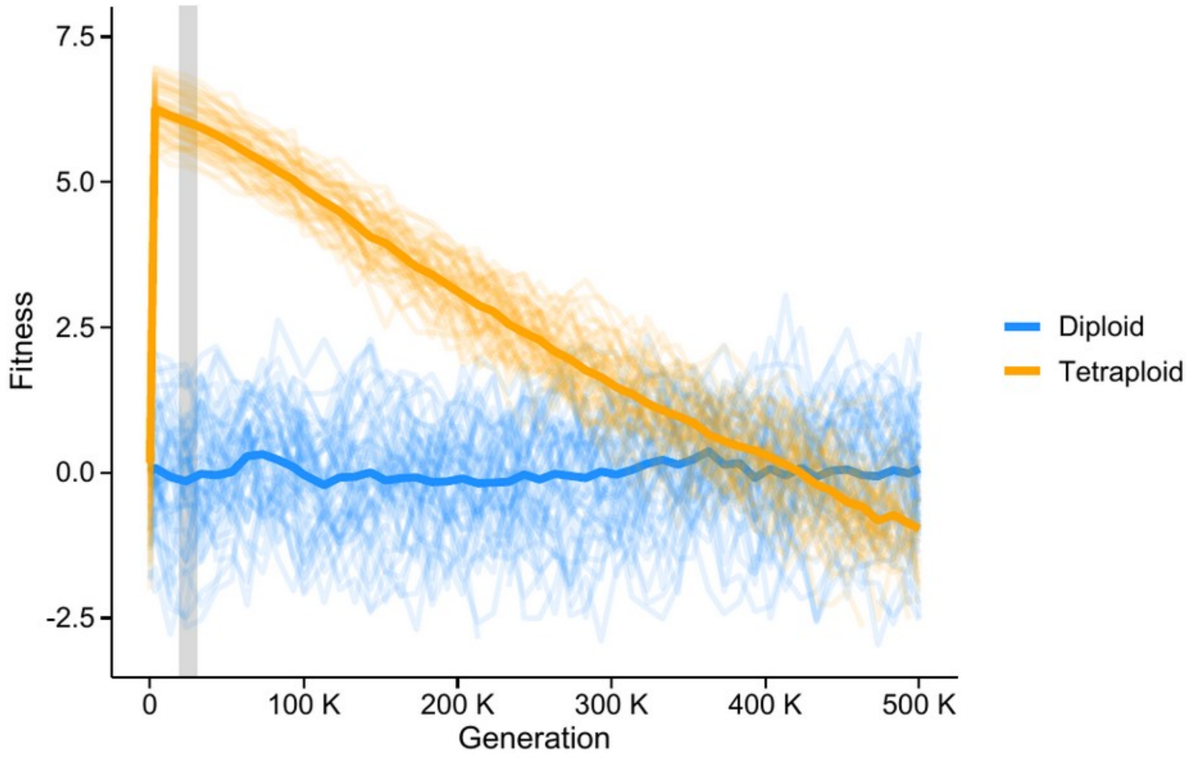


**Figure S3:** Simulated fitness over 500K generations after WGD in a scenario without initial tetraploid bottleneck.

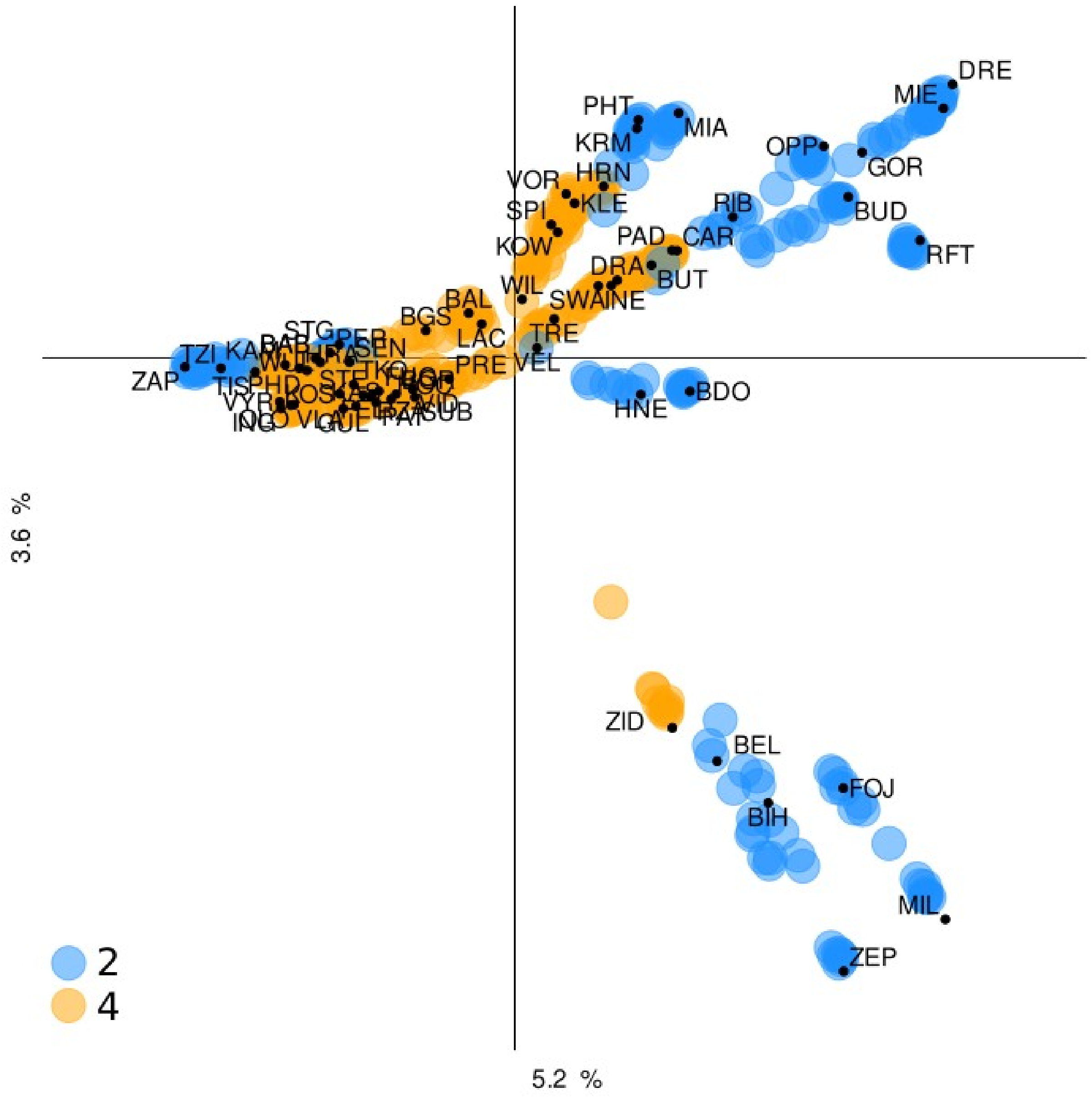


**Figure S4:** Principal component analysis of genetic relationship of all short-read sequenced individuals. The PCA is based on 92,632 4-fold degenerate SNPs.

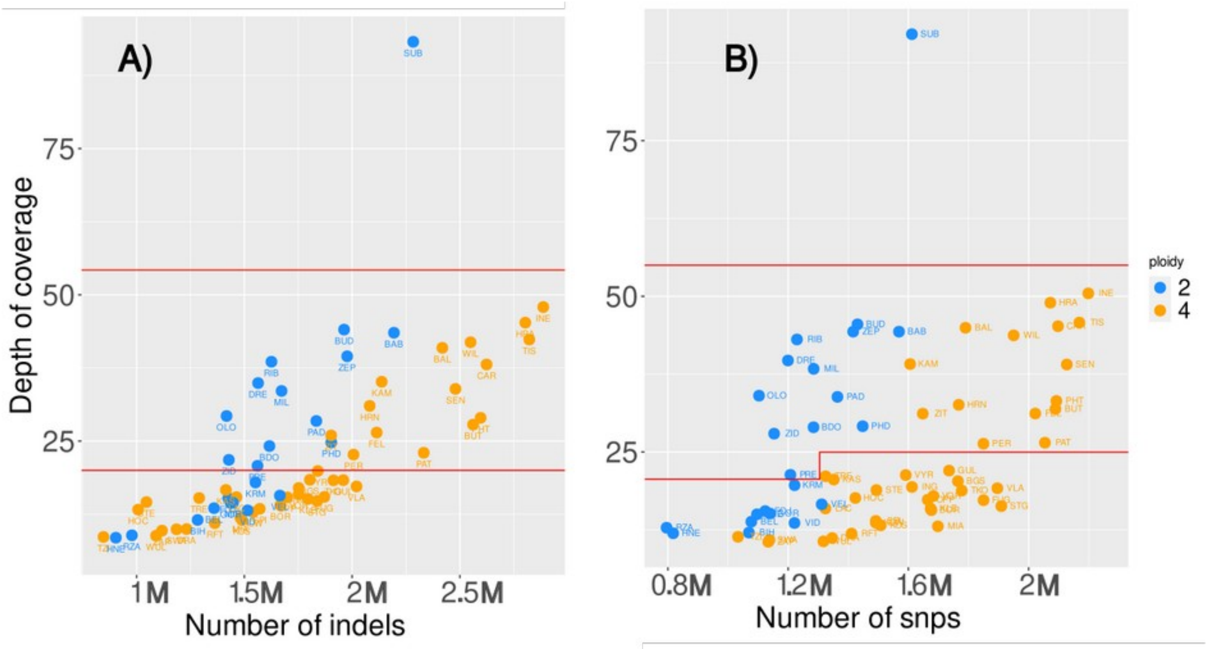


**Figure S5:** Mean depth of coverage of A) indels and B) SNPs per population (dot) and their total sum per population downsampled to 6 individuals. Area between red lines indicate a core subset of populations that was kept the same for both markers.

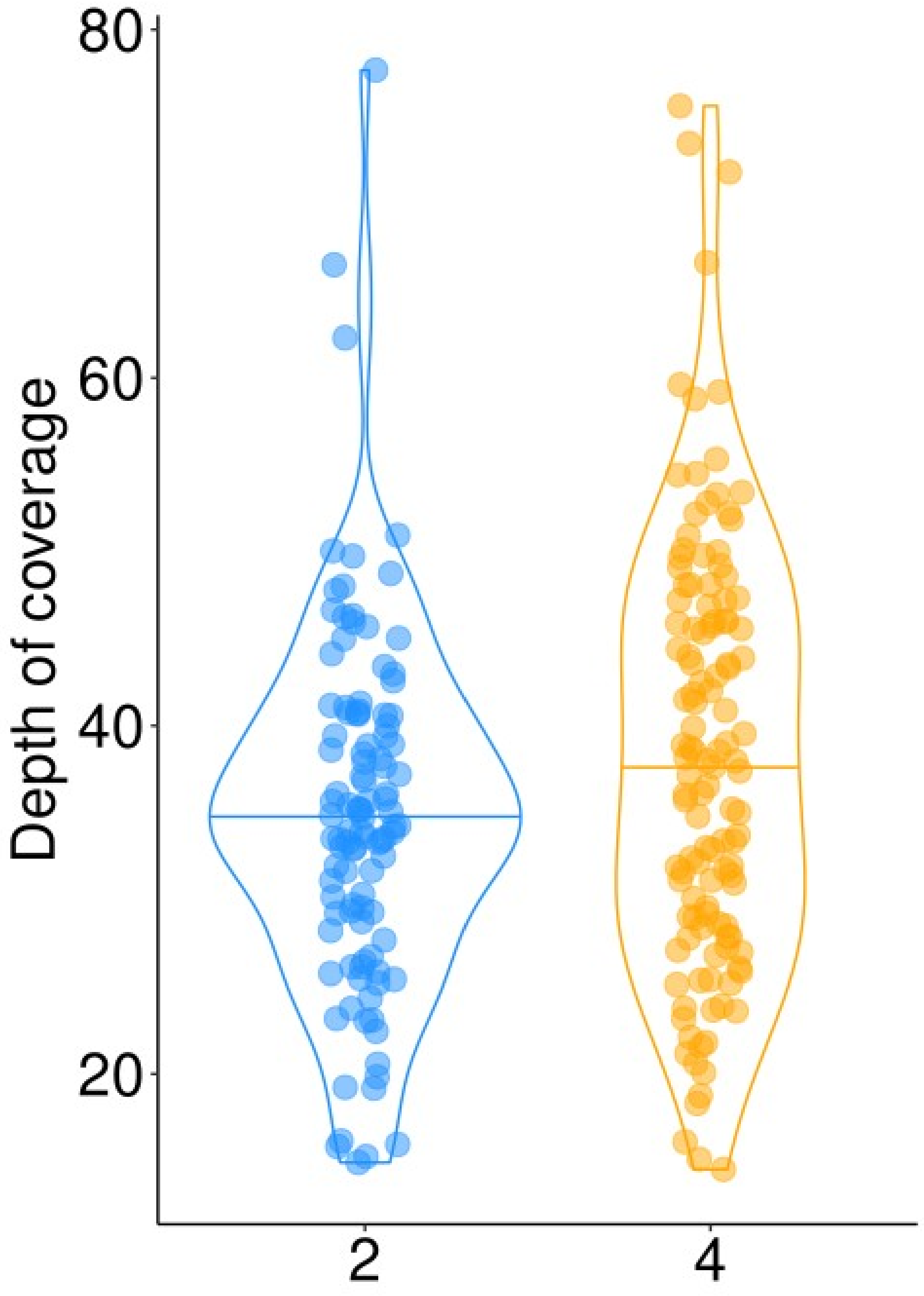


**Figure S6:** Depth of coverage of four-fold degenerate SNPs per individual in the core dataset does not differ between ploidy (p = 0.548, Wilcoxon test).

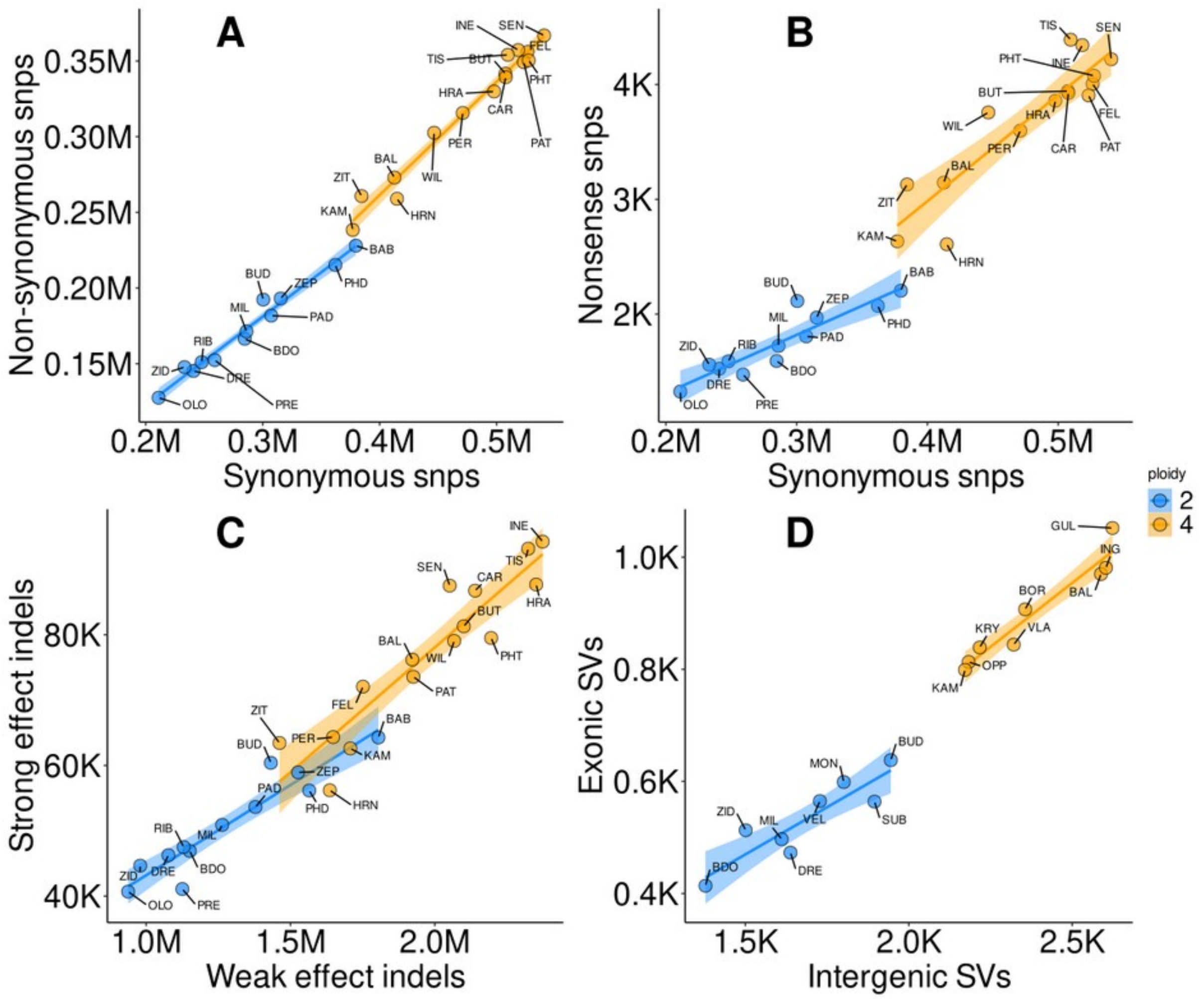


**Figure S7:** Fully annotated version of the Figure 4 with labels of individual populations. Relationship between putatively neutral (horizontal axis) and deleterious (vertical axis) SNPs, insertion-deletion (indel) and structural variants (SVs) in diploid (blue) and tetraploid (orange) populations. Population data (SNP and indels) were downsampled to 6 individuals, SVs were detected using one long-read sequenced individual per population. Tetraploids show not only consistently more variants of both categories, but also proportionally higher accumulation of deleterious variants (slope, β), an indicator of weakened efficiency of purifying selection. Note that β was calculated for scaled and normalised data to allow direct comparison between site categories. A) Number of minor non-synonymous SNPs as a function of number of minor synonymous SNPs per population. B) Number of minor nonsense SNPs (inserting premature stop-codons) as a function of number of minor synonymous SNPs per population downsampled to 6 individuals.). C) Number of minor strong effect indels as a function of number of minor weak effect indels per population downsampled to 6 individuals. D) Total number of exonic structural variants (SVs) as a function of total number of intergenic SVs per individual.

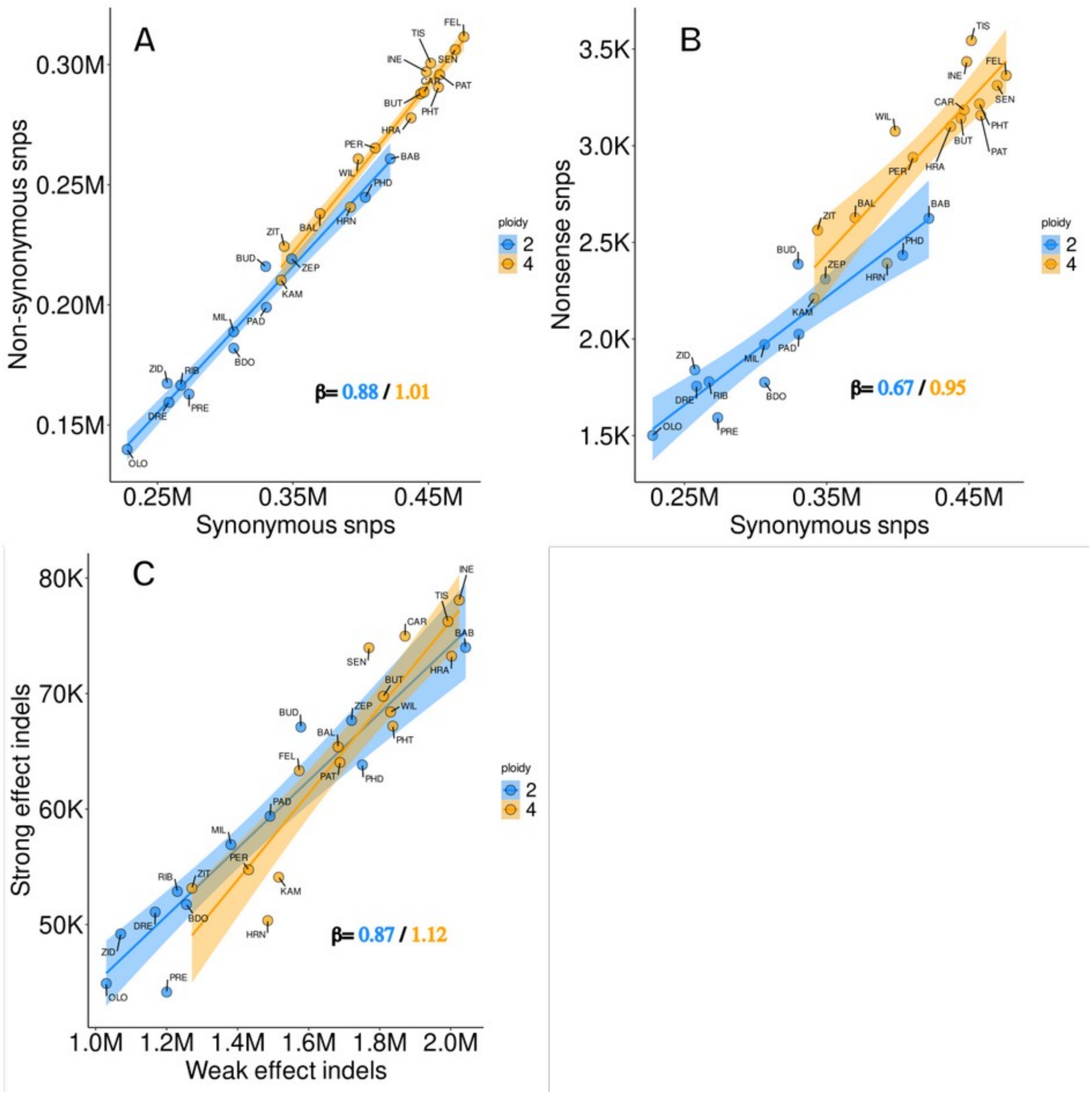

**Figure S8:** Indices of load inferred as number of variant sites normalised to the same number of sampled chromosomes per ploidy. Four tetraploids and 8 diploid individuals per population were used to unify the number of sampled chromosomes between cytotypes to 16. A) Number of non-synonymous vs. synonymous SNPs, B) Number of nonsense vs. synonymous SNPs, C) Number of strong effect indels vs. weak effect indels. Interaction of Ploidy and Synonymous alleles is significant according to likelihood ratio test comparing a null model with only a direct effect of each factor with a model also including their interaction by LRT test in all cases (Table S4).

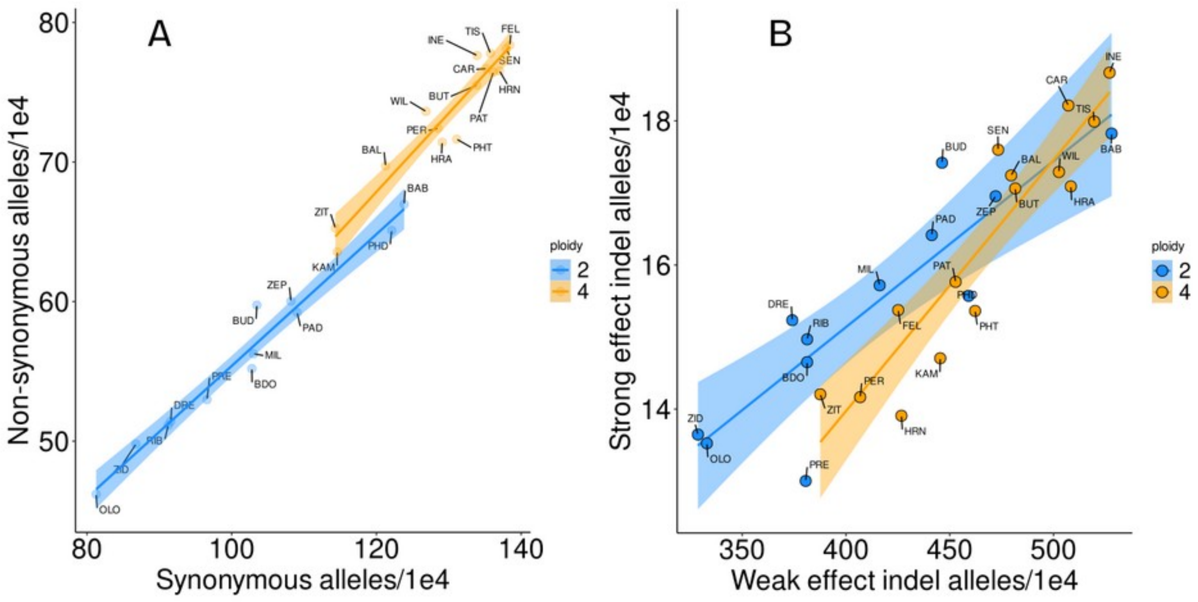


**Figure S9:** Total number of *alleles* from normalised dataset sampling 16 chromosomes per each population independent of ploidy. The total number of alleles reflects the total load. A) Normalised number of nonsynonymous vs. synonymous SNP alleles. B) Normalised number of strong effect vs. weak effect indel alleles. Interaction of of Ploidy and Synonymous alleles is significant according to likelihood ratio test comparing a null model with only a direct effect of each factor with a model also including their interaction by LRT test in both cases (p<0.000).

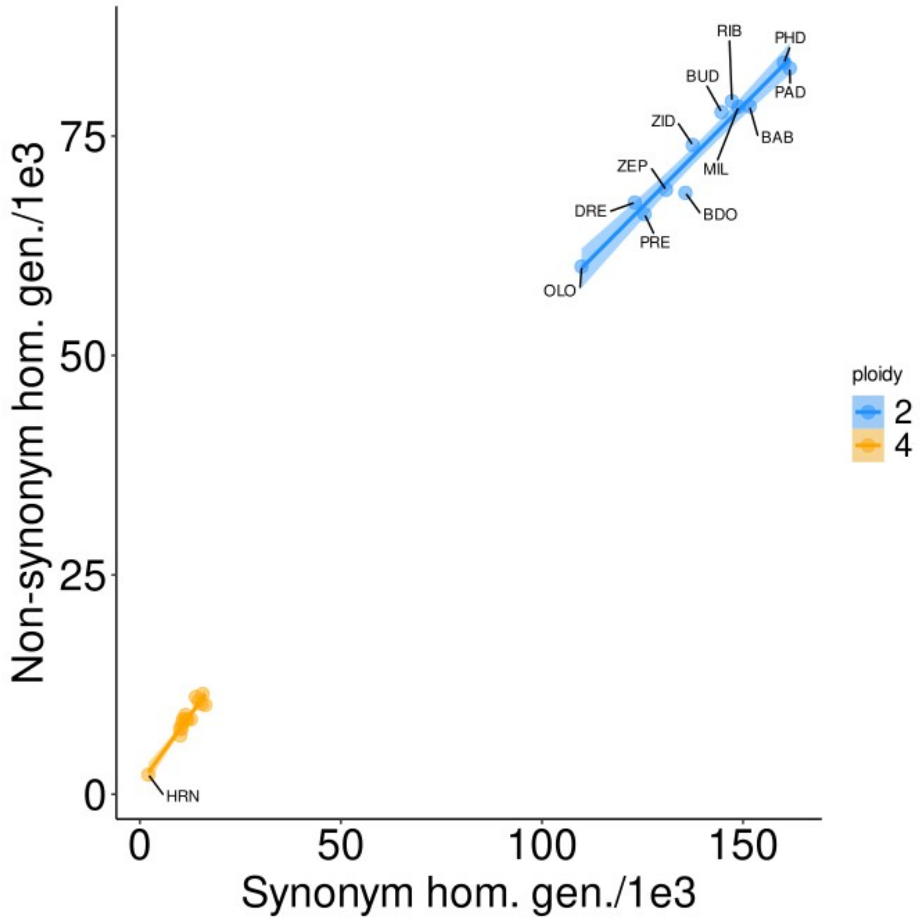


**Figure S10:** Number of *homozygous genotypes* carrying non-synonymous vs. synonymous SNPs used as a proxy for realised load. Four tetraploids and 8 diploid individuals per population were used to unify the number of sampled chromosomes between cytotypes to 16.
